# HumanIslets: An integrated platform for human islet data access and analysis

**DOI:** 10.1101/2024.06.19.599613

**Authors:** Jessica D. Ewald, Yao Lu, Cara E. Ellis, Jessica Worton, Jelena Kolic, Shugo Sasaki, Dahai Zhang, Theodore dos Santos, Aliya F. Spigelman, Austin Bautista, Xiao-Qing Dai, James G. Lyon, Nancy P. Smith, Jordan M. Wong, Varsha Rajesh, Han Sun, Seth A. Sharp, Jason C. Rogalski, Renata Moravcova, Haoning H Cen, Jocelyn E. Manning Fox, HI-DAS Consortium, Ella Atlas, Jennifer E. Bruin, Erin E. Mulvihill, C. Bruce Verchere, Leonard J. Foster, Anna L. Gloyn, James D. Johnson, Andrew R. Pepper, Francis C. Lynn, Jianguo Xia, Patrick E. MacDonald

## Abstract

Comprehensive molecular and cellular phenotyping of human islets can enable deep mechanistic insights for diabetes research. We established the Human Islet Data Analysis and Sharing (HI-DAS) consortium to advance goals in accessibility, usability, and integration of data from human islets isolated from donors with and without diabetes at the Alberta Diabetes Institute (ADI) IsletCore. Here we introduce <u>HumanIslets.com</u>, an open resource for the research community. This platform, which presently includes data on 547 human islet donors, allows users to access linked datasets describing molecular profiles, islet function and donor phenotypes, and to perform various statistical and functional analyses at the donor, islet and single-cell levels. As an example of the analytic capacity of this resource we show a dissociation between cell culture effects on transcript and protein expression, and an approach to correct for exocrine contamination found in hand-picked islets. Finally, we provide an example workflow and visualization that highlights links between type 2 diabetes status, SERCA3b Ca^2+^-ATPase levels at the transcript and protein level, insulin secretion and islet cell phenotypes. <u>HumanIslets.com</u> provides a growing and adaptable set of resources and tools to support the metabolism and diabetes research community.

## INTRODUCTION

Four decades following the development of methods for large-scale isolation of human pancreatic islets,^1–3^ the use of cadaveric islets has become a staple of diabetes research.^4–7^ As biomedical datasets continue to grow in size, there has been increased emphasis on improving the findability, accessibility, interoperability, and reusability (FAIR) of published data.^8–10^ This is highly relevant to human islet research^6^ since these are increasingly used as a comparator in the development of stem cell-derived products for transplantation,^11–13^ in large genomics studies at the islet^14,15^ and single-cell level,^16,17^ and in studies that seek to understand conventional type 1 diabetes (T1D) and type 2 diabetes (T2D).^4^ Furthermore, since islet tissue is not readily accessible it is particularly important to maximize the usage of existing datasets.

Several initiatives provide data resources and analysis pipelines for the exploration of human islet data, with differing degrees of openness. Some focus on deep human pancreas phenotyping and data curation,^7,18–21^ while others provide analysis tools for linking hormone production and gene expression,^22^ single-cell islet transcript visualization,^23,24^ and exploring the relationship between diabetes genetics and gene regulation.^25^ While existing resources are valuable, there is an unmet need both for user-friendly analysis pipelines that are accessible to the general islet and diabetes research community, including non-specialists, and for improved access to analysis pipelines and minimally processed data for more customized analyses.

Additionally, since there is often low concordance in findings from different datasets and analyses,^4^ a better understanding of how donor factors, technical aspects of organ processing, and cell culture impact these data is crucial for appropriate interpretation of results. Proper documentation, reporting, and integration of this information into transparent data collection and analysis pipelines can make important contributions to understanding islet data and diabetes research. We established the HumanIslets Data Analysis & Sharing (HI-DAS) Consortium to address these important gaps in the human islet research ecosystem.

The Alberta Diabetes Institute (ADI) IsletCore (www.isletcore.ca) isolates and distributes human islets specifically for research. Together with HI-DAS we present an online resource (<u>HumanIslets.com</u>) for the exploration, analysis, and download of data collected from human research islet preparations from organ donors with and without diabetes. This platform includes data (547 donors as of June 2024) and methods for analysis of bulk islet transcriptomics, bulk islet proteomics, and single-cell transcriptomics linked to measures of islet functions such as insulin secretion, oxygen consumption, electrophysiology, pancreas processing, and cell culture metadata. The <u>HumanIslets.com</u> platform offers intuitive support to directly analyze these data and metadata, including: ***i)*** exploration at the donor, islet and single-cell levels; ***ii)*** multi-omic analysis of genes/proteins and pathways controlled for common factors that may influence results; and ***iii)*** download of data, results and analysis methods for transparent and reproducible re-analysis. Furthermore, we demonstrate the impact of correcting for culture time and exocrine contamination following cell type composition deconvolution using bulk proteomics data. This platform is under continuous growth as new donors and additional data types are added and is freely available for the diabetes and islet research community.

## RESULTS

The <u>HumanIslets.com</u> tool is a web-based platform for accessing, analyzing, and visualizing phenotyping data collected from human research islets isolated at the ADI IsletCore. The underlying database includes technical metadata on the islet isolation and cell culture process; multi-omics; functional data; and associated donor metadata that describes basic, de-identified demographic parameters (**Figure 1**). The tool is designed to be easily expanded with new data types, and regularly updated as data are collected from new donors.

**Figure 1.**
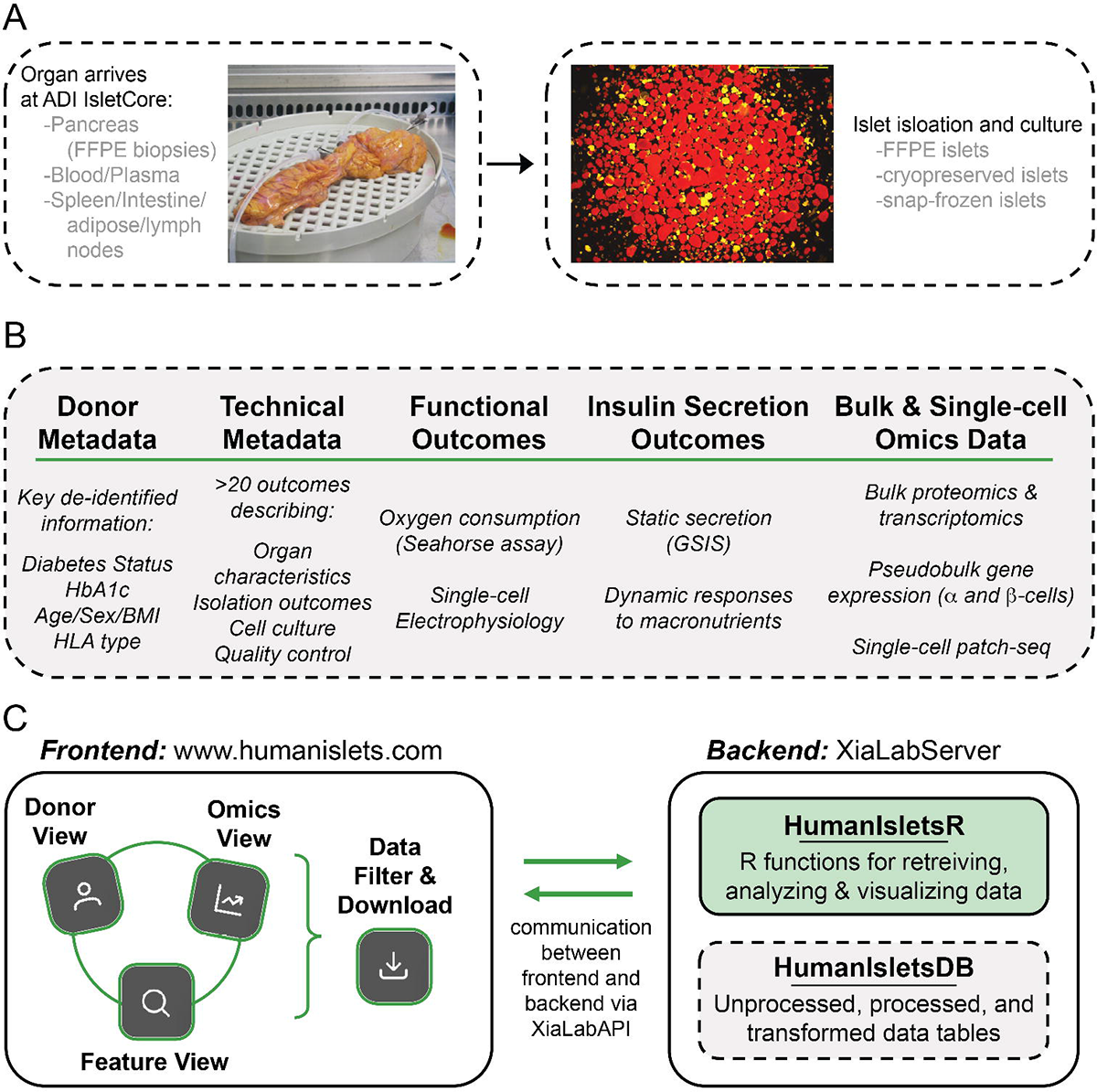
A human research islet phenotyping, sample and data collection, and data analysis and visualization pipeline. **A)** Human research pancreas received by the Alberta Diabetes Institute (ADI) IsletCore is processed for sample collection (grey) and islet isolation (FFPE – formalin fixed paraffin embedded). **B)** Metadata and data are collected on the organ donor, organ and tissue processing, islet and cellular function, and molecular profiling via several omics platforms. **C)** The *HumanIslets* tool comprises both a backend server for data storage and computation, and a user-friendly frontend for interactive data analysis and visualization.

### Database contents

The *HumanIslets* platform presently contains data from 547 donors. These are 41.1% female, range in age from 10-weeks to 84 years, and include donors with no diabetes (n=355, 65%), pre-diabetes (n=99, 18%, defined as HbA1c of 5.7-6.5% and no clinical diagnosis of T2D), T2D (n=78, 14.3%, defined as a reported diagnosis of T2D or HbA1c>6.5%), and T1D (n=15, 2.7%, defined as a reported diagnosis of T1D, typically with validation by autoantibody screening, genetic risk scores, and/or biopsy immunostaining) (**Figure 2**). The donor metadata also contain technical details on pancreas characteristics and processing, islet isolation, and cell culture outcomes. There are extensive functional phenotyping data including static insulin secretion at various glucose concentrations, dynamic insulin secretion in response to glucose and other macronutrients, islet oxygen consumption in response to glucose and mitochondrial modulators, and single-cell electrophysiological outcomes for both α- and β-cells. Certain data are available for all donors, while others have been collected from more recent subsets or are not yet available (**Figure 2E**). In total, there are 74 distinct donor, technical, and functional outcomes summarized at the donor level, providing a rich context within which to analyze and interpret islet multi-omics data from the same donors (**Suppl Table 1**).

**Figure 2.**
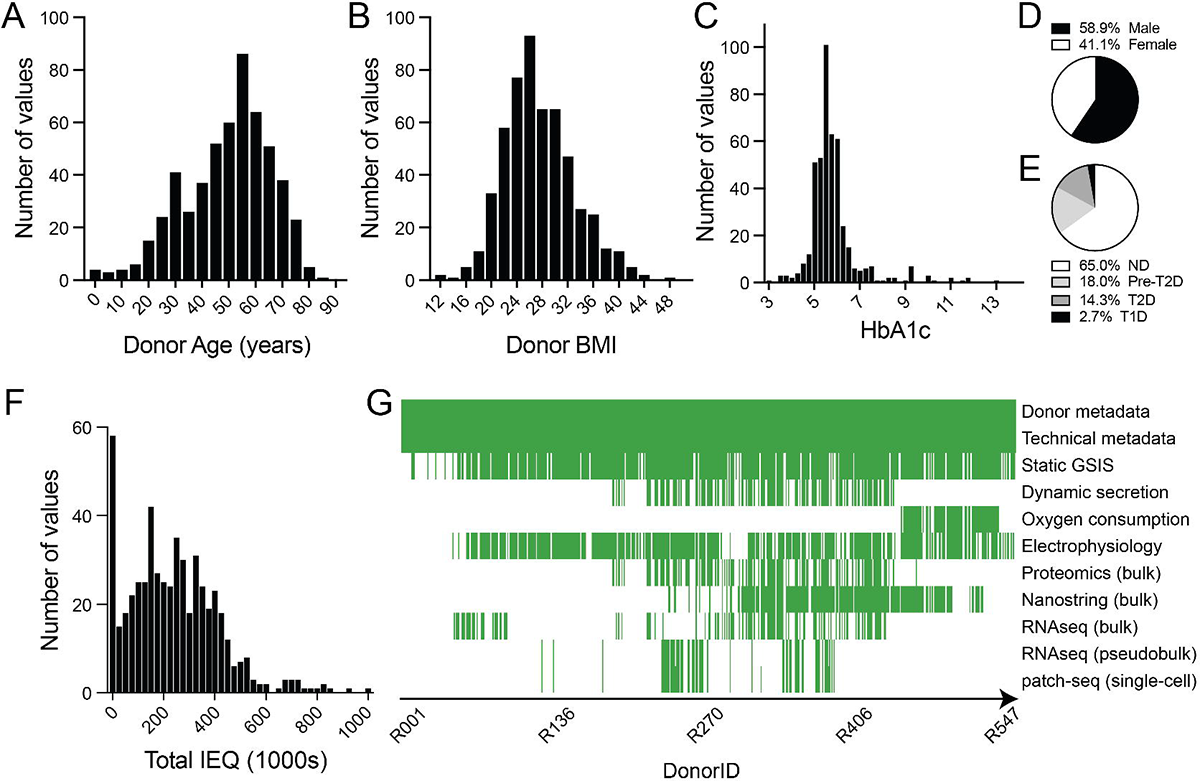
Profiles of donors, islets and data types within the database. **A-E)** Distribution of donor age (A), BMI (B), HbA1c (C), sex (D) and corrected diabetes status (E) across the 540 donors currently in the database. **F)** Distribution of islet isolation outcomes across the 540 donors currently in the database. **G)** Heatmap showing the availability of metadata and data types within the database and for download.

Bulk transcriptomics, bulk proteomics, and single-cell transcriptomics were collected from isolated islets of subsets of the donors. Two different platforms were used for the bulk transcriptomics: including RNA-seq (*n*=117 donors; whole transcriptome) and NanoString (*n*=190 donors; 156 genes). The single-cell transcriptomic data are available for analysis at the single-cell or donor level. At the single-cell level, transcriptomes are linked with nine single-cell electrophysiology outcomes (Patch-seq data; *n*=∼4300 cells collected from 96 donors) sub-setted by cell type (α-, β-cells) and glucose concentration (1, 5, and 10mM). For donor-level outcomes (ie. age, diabetes status), the data are summarized as a single α- and β-cell pseudobulk profile for each donor. Analysis of single-cell data with respect to donor-level outcomes like diabetes status can lead to artificially low p-values when each cell is treated as an independent replicate by not incorporating donor IDs into the statistical analysis (known as ‘pseudoreplication’).^26^ Two main solutions have been proposed: analyzing pseudobulk profiles with respect to donor-level outcomes,^27,28^ or using linear mixed models with donor incorporated as a random effect.^26^ Given there is no consensus on the best approach,^29^ we use the pseudobulk profiles method to support interactive visual exploration. Users can download the data and corresponding analysis script to perform mixed model analysis, if desired.

### Platform design

The <u>HumanIslets.com</u> platform was developed using a design-thinking approach^30^ to focus features on addressing the needs of the islet and diabetes research community. Within the HI-DAS Consortium, tool developers were in continual contact with islet biologists across career stages and with broad expertise across domains of islet biology and diabetes, enabling feedback on prototypes followed by multiple iterations of each feature. This process revealed a clear desire for support for simple queries and common analyses of well-organized and documented data via an interactive web interface, coupled with the ability to easily subset and export minimally processed data for more customized and sophisticated offline analyses. This focused the tool design on intuitive workflows using well established and widely understood statistical methods, providing clear documentation of all steps, and presenting results with familiar visualizations. The platform is designed to allow researchers to query statistical associations within the data by omics type (*Omics View*), by individual omics or metadata feature (*Feature View*), or by donor (*Donor View*) (**Figure 1C**). A visual walkthrough of the functions described below is presented in the **Supplementary Document 1**.

#### Omics View

allows researchers to find omics features and pathways associated with any of the 74 distinct donor, technical, and functional outcomes. The interface allows users to select an omics type, a metadata variable of interest, and any covariates that should be adjusted for.

Researchers can perform the analysis on a subset of donors by applying different filters and can specify the p-value significance threshold. The association analysis results are displayed in an interactive volcano plot coupled with a detailed result table. Clicking on a point (in volcano plot) or result row (in table) will display a scatterplot or violin plot for the association/analysis of interest which can be exported in PNG, PDF, or SVG format. Within these, the donor-level data points are interactive - clicking them will open the ‘*Donor View*’ page query results for that donor, allowing researchers to quickly investigate outliers and to assess whether they are driven by technical or biological factors. The result table provides raw and adjusted p-values as well as estimates for cell type associations (*Associated Proportions*) based on deconvolution analysis of the bulk proteomics data. Links in this table take users to the relevant gene page at the National Center for Biotechnology Information (‘NCBI*’*) or Type 2 Diabetes Knowledge Portal (‘T2DKP*’*),^31^ or generate a figure summarizing the cell-type specific gene expression across 14 cell types from 109 donors without diabetes, representing more than 140,000 individual cells (‘*SC’*).

Clicking on the feature name (gene/protein) will link directly to the *Feature View* page (see below) for that hit. Finally, using *Pathway Analysis*, data can be analyzed using either overrepresentation analysis (ORA) or gene set enrichment analysis (GSEA). Statistically significant pathways are displayed in an interactive ridgeline plot that allows comparison of relative positive and negative metadata associations across different pathways. Clicking on a pathway of interest will generate heatmaps of protein or gene expression for all features in a pathway. Pathway analysis results can also be viewed in table form (Results Table). This allows researchers to quickly perform and visualize standard differential expression, association, and pathway analyses, compare global signals across omics layers, and explore the influence of different covariates on the results. When users perform an analysis in the ‘Omics View’ page, the list of donors with completely paired sets of the omics data and metadata used in the analysis is made available for download. All associations and pathway results, along with the R scripts that generated them, can be downloaded directly via the ‘Download Results’ and ‘Save Analysis’ buttons, respectively.

#### Feature View

consists of a search bar and database of pre-computed statistical associations describing over 3.7 million relationships, including pairwise combinations of all omics features and metadata variables (omics-metadata), while adjusting for influential technical and biological covariates (sex, age, BMI, culture time). Users can enter the name of any variable within the *HumanIslets* database to retrieve its associations with all other outcomes. If a gene name or symbol is searched, significant associations for both the protein and mRNA gene products will be included in the search results, which can be subsequently filtered by gene/protein symbol, metadata variable, or omics type. This page was designed for researchers who have a particular gene or outcome of interest, and would like to find out which proteins, mRNAs, or metadata variables are most correlated with it.

#### Donor View

allows users to query a specific donor by ADI IsletCore DonorID or RRID, and returns that donor’s metadata and functional outcomes, highlighted within the context of all other donors in the database. For example, searching donor ‘R421’ displays a histogram of HbA1c values from all donors, with donor R421’s value of 5.4% highlighted with the percentile rank (0.38, or 38^th^ percentile), and the same for age and BMI. The Donor View page has nine different tabs, each presenting a different category of data. Clicking through these will show, in addition to donor information: the properties and images of the islet isolation itself; insulin content results and static insulin secretion across glucose concentrations; dynamic insulin secretion responses to glucose, leucine and oleate/palmitate; oxygen consumption assessed via Seahorse assay; and electrophysiological profiles of α-cells and β-cells. Each of these data are shown in relation to the entire dataset, allowing researchers to understand how comparable a given donor is to the rest of the population represented within the database: this is a common question when a donor appears as an outlier in a particular analysis or when a research group receives islet samples from the ADI IsletCore from a small number of donors. Finally, the last tab presents a summary of omics data available in the tool for that donor.

#### Data Download

allows researchers to download all data from the *HumanIslets* database using extensive filtering options to subset donors using common or advanced criteria, by uploading a list of donor IDs, or by dataset availability. For example, a researcher could filter the list to include only female donors with both insulin secretion and bulk RNA-seq data. The omics data are available for download in their unprocessed and processed forms. The unprocessed data refer to the original omics data table; for example, intensity values for proteomics data and counts for RNA-seq-based gene expression data. The processed omics data have undergone batch correction, missing value imputation, filtering, and normalization (see Methods for more details). Raw omics data (ie. proteomics MS spectra, sequencing FASTQ files) are not stored in the <u>HumanIslets.com</u> tool but are accessible in public repositories (links provided under ‘Documentation’ and in Data Accessibility below). The final data tables can be exported in either CSV, plain text, Excel format, or as an R object, together with a data dictionary outlining units and any formulas used to calculate each variable. Details on data collection and processing are available in the ‘Documentation’ (**Suppl Document 1**) link at the top of the web page.

### Transparency, reproducibility, and maintainability

The <u>HumanIslets.com</u> platform comes with extensive documentation on how each dataset was generated and the precise definition of each variable. A brief description is given for all statistical methods, and how they are combined into pipelines to generate the results. The HumanIsletsR Github repository (https://github.com/xia-lab/HumanIsletsR) provides all R functions used to perform analyses within the tool, and all R scripts used to process each dataset. The processing scripts are designed to dynamically pull together data from different sources as various datasets are expanded and save the data at different stages of processing for download, analysis, and visualization within the tool. Some steps, such as outlier detection and omics data normalization, depend on statistics computed from all samples and therefore produce slightly different processed values as new donors are added (or in comparison to the original published analyses of individual datasets). This presents a reproducibility problem, as the same analysis on the omics View page will produce slightly different results after the data are updated, even if the donors are filtered to include the same list of IDs. Therefore, the “*Download Analysis*” feature was added to the tool, which allows researchers to download a time-stamped copy of the processed omics data, paired metadata outcomes, and R functions, allowing them to fully reproduce the analysis offline.

### Important technical and biological associations

The omics data were analyzed with respect to each metadata variable to understand the relative signal strength associated with different functional outcomes, and to identify which donor and technical covariates were influential across multiple omics layers. These results informed the linear model used to generate the pre-computed association results in the ‘*Feature Search*’ page. Here we highlight overarching trends across the entire database and focus on technical covariates that are often overlooked.

The number of omics features statistically significantly associated with eight key donor and technical metadata are presented in **Table 1**, along with the dimensions of each omics data and whether there were significant batch effects present, details that are important to consider when interpreting the number of significant features across modalities. ‘Culture time’ and ‘percent purity’ are two metadata categories with the most significantly associated features in multiple omics datasets (**Table 1**). Both are technical parameters and should be carefully considered in the analysis and interpretation of data. The metadata comparison with the next highest number of significantly associated features across omics layers was T2D-ND. HbA1c (%) was associated with a high number of features in the proteomics data only. Age and sex also had a moderate number of associated features across multiple omics platforms. Most mRNAs and proteins significantly associated with sex are encoded by genes found on the X- and Y chromosomes, but do not show up in the NanoString data because they were not included in that gene panel; genes in the NanoString panel were mainly selected for their relevance to islet function and development. This handful of explicitly sex chromosome-linked features is strongly associated (FDR<10^-10^) with sex, while most other mRNAs and proteins have no discernible difference between male and female donors.

**Table 1.** Number of omics features significantly associated with key metadata (adj. P-value < 0.05, FDR method). Table cells with an “N/A” indicate that the analysis was not performed because there were fewer than 10 donors with omics profiles per group for categorical variables, or fewer than 10 donors overall for continuous variables. Note that HbA1c data are not available for all donors and there were a few donors with T1D, thus the sample size is slightly less than the number of omics profiles for some of the T2D-ND and HbA1c analyses.

|  | **Bulk RNA-seq** | **Nanostring** | **Proteomics** | **Pseudobulk RNA-seq ($\alpha$)** | **Pseudobulk RNA-seq ($\beta$)** |
| --- | --- | --- | --- | --- | --- |
| **Features** | 14 314 | 156 | 6138 | 16 874 | 14 950 |
| **Donor profiles** | 117 | 190 | 134 | 52 | 30 |
| **Significant batch effects** | Yes | Yes | No | No | No |
| **Key diabetes outcomes** |  |  |  |  |  |
| **T2D-ND** | 418 | 51 | 1519 | 0 | N/A |
| **HbA1c (%)** | 5 | 94 | 781 | 0 | 5 |
| **Selected clinical covariates** |  |  |  |  |  |
| **Age (years)** | 105 | 39 | 46 | 0 | 0 |
| **Sex** | 27 | 0 | 20 | 12 | 8 |
| **BMI** | 0 | 1 | 0 | 0 | 0 |
| **Selected technical covariates** |  |  |  |  |  |
| **Cold ischemic time (h)** | 2 | 0 | 2 | 0 | 0 |
| **Purity (%)** | 2245 | 98 | 2414 | 1 | 0 |
| **Culture time (h)** | 7547 | 30 | 1 | 0 | 0 |

There were many fewer, or no, pseudobulk mRNA features associated with any metadata variable except for sex, which makes sense given the lower sample size and because these profiles are derived from single-cell data that has greater noise, lower sequencing depth, and higher drop-out rates compared to bulk data. However, while the adjusted p-values may not be statistically significant at the feature level, analyzing the pseudobulk association results at the pathway level with GSEA produces statistically significant results that are consistent with known biology. For example, comparing pseudobulk mRNA expression in α-cells from donors with and without T1D highlights the upregulated Reactome pathway ‘Antigen Presentation: Folding, assembly and peptide loading of class I MHC’ as shown by others.^32^ An expanded table with the full association results for all metadata variables is available in the Supplementary Information (**Suppl Table 2**).

### Culture time has less impact on protein expression than transcript expression

The distribution of culture time values is multimodal with each peak spaced roughly 24 hours apart (**Figure 3A**) because islets were typically removed from culture during business hours. Consistent with what we recently observed,^33^ a large proportion of features were associated with culture time in the bulk RNA-seq (53%; **Figure 3B**) and NanoString (32%) data, but not the bulk proteomics data (∼0.01%; **Figure 3C**). Gene products tend to have consistent relationships across the bulk RNA-seq and NanoString data, but not with the proteomics data. For example, *FGF2* (fibroblast growth factor 2) is positively associated with culture time in both the bulk RNA-seq (FDR=5.3e-9, **Figure 3D**) and NanoString (FDR=4.1e-6, **Figure 3E**) data, but not in the proteomics data (FDR=0.99, **Figure 3F**). Pathway analysis of the bulk RNA-seq data with GSEA (features ranked by T-statistic) using the KEGG, Reactome, and Gene Ontology Biological Process libraries consistently identified proteasome, spliceosome, and ribosome-related gene sets as the most significantly enriched with culture time, along with evidence for altered immune pathways, cell cycle pathways, oxidative phosphorylation, and fatty acid metabolism (**Figure 3G; Suppl Table 3**). Consistent with the idea that these changes in gene expression have not (yet) translated into alterations in protein expression and functionality, analysis of insulin secretion from these donors show no impact of culture time on stimulation index within this time frame.^34^

**Figure 3.**
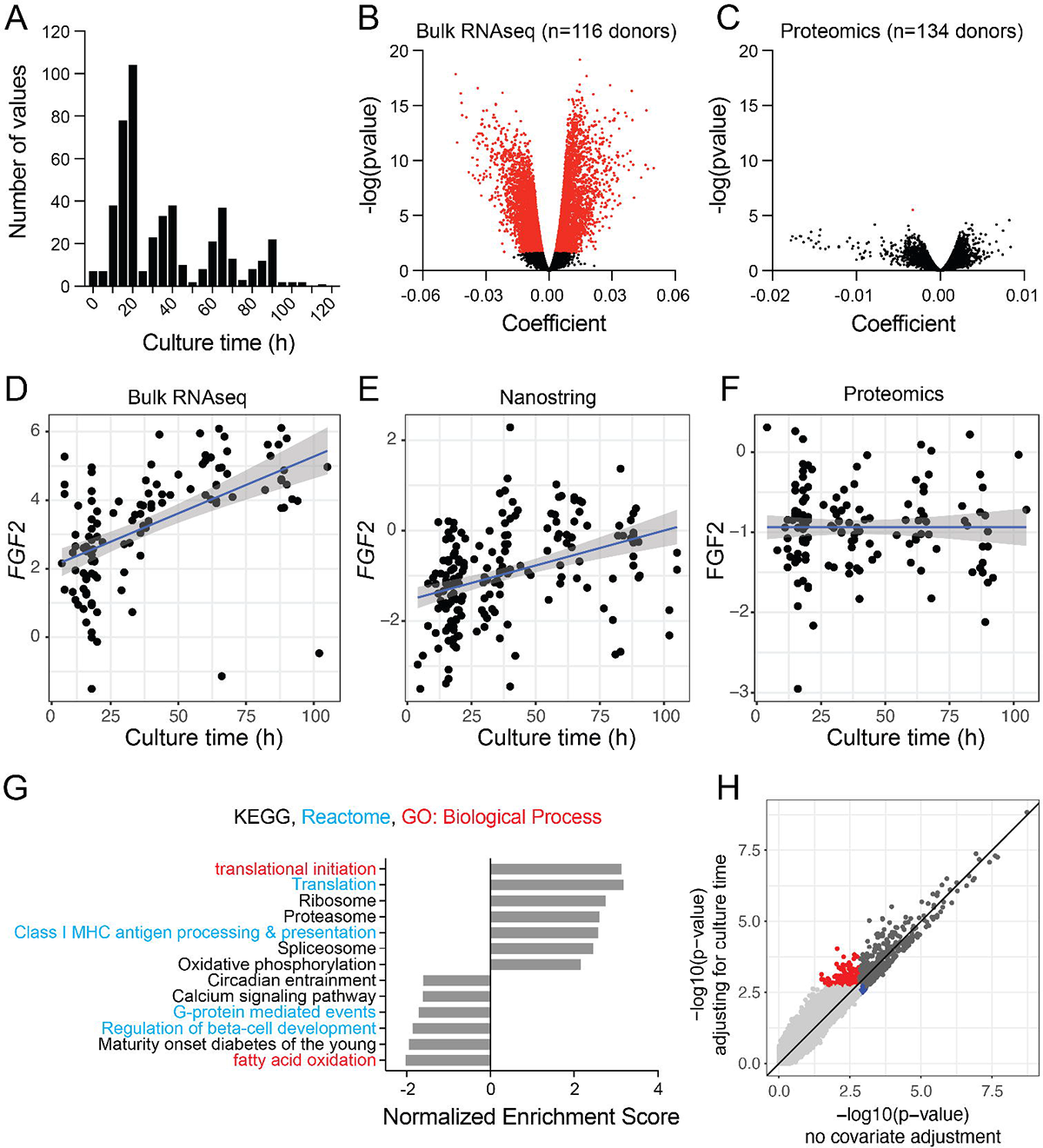
Effect of culture time on transcript and protein expression, and the impact of including ‘culture time’ as covariate in analysis. **A)** The time in culture following research islet isolation and prior to sample collection or shipment. Times are clustered due to collection and shipment during business hours (usually in the morning). **B-C)** Effect of culture time on transcript (B) and protein (C) expression in human islets. Adjusted p-value <0.05 indicated in red. **D-F)** Impact of culture time on expression of *FGF* measured by RNA-seq (D), NanoString (E), or FGF measured by proteomics (F). **G)** Gene set enrichment analysis using T-statistic as weighting against KEGG (black), Reactome (blue), and GO: Biological Process (red) genesets. Selected significant pathways (FDR<0.05) are shown. See also **Suppl Table X**. **H)** Impact of adjusting for culture time on gene-level p-values for the association of bulk RNA-seq data and diabetes status (T2D vs. None). Transcripts were either significant (FDR<0.05) both with and without adjusting for culture time (dark grey), only when adjusting for culture time (red), only when not adjusting for culture time (blue), or not significant in either scenario (light grey).

### Control for culture time in data analysis

Culture time is largely determined by the time of day and week that the ADI IsletCore receives donor organs, and is not significantly associated with any donor metadata, technical metadata other than other measures of processing, or functional outcomes (correlation for continuous metadata, ANOVA for discrete metadata, FDR<0.05). Users can control for culture time by including ‘culture time’ (under ‘Cell Culture Outcomes’) as a covariate when performing analysis on the ‘*Omics Analysis*’ page. Accounting for culture time when analyzing the bulk transcriptomics data improves the strength of the relationships between gene expression and nearly every other variable, resulting in decreased p-values for associated transcripts and a higher number of transcripts that pass the significance threshold (**Figure 3H**). For example, the number of transcripts with significant differential expression between donors with and without T2D increases from 418 to 542 (RNA-seq) and from 25 to 30 (NanoString) when culture time is included as a covariate.

### Impacts of cell type composition

The purity of the islet preparation (percent purity) is estimated by the isolation research technician,^35,36^ leading to more frequent instances of whole numbers (**Figure 4A**). Purity is significantly associated with many transcripts and proteins in the bulk omics data (**Figure 4B**). This initially seems surprising since the islets were hand-picked to nearly 100% purity prior to acquiring these omics data. However, purity is negatively correlated with the ‘percentage trapped’ islets (islets embedded within acinar fragments, r = −0.28, p-value = 1.88e-11, Pearson’s correlation) and is significantly lower in donors with diabetes compared to those without (two-sided t-test, p-value = 1.39e-7 for T2D versus no diabetes and p-value = 3.93e-17 for T1D versus no diabetes). Single-cell gene expression distributions show that bulk RNA-seq and proteomics features with the strongest positive and negative associations with purity have biased expression in endocrine and non-endocrine cell types respectively (**Suppl Figures 1,2**), suggesting that their relationship with purity is explained by differential amounts of non-endocrine cells that get trapped on islet surfaces and that persist after hand picking. Thus, lower initial islet purity could be associated with more acinar tissue (even in hand-picked islets) as well as islet dysfunction. It may therefore be difficult to distinguish features that are direct reflections of lower purity preparations from those with perturbed biological function. To address this, we used the abundance of known endocrine marker proteins to estimate the relative amounts of endocrine (β-, α-, δ-, and pancreatic polypeptide (PP) cells) and non-endocrine cells in the samples with proteomics analysis (**Figure 4D**). The estimated non-endocrine proportion is negatively correlated with purity (*r* = −0.54, p-value = 1.13e-11, Pearson correlation), and has the highest number of significantly associated features across both the bulk proteomics and transcriptomics data (**Table 2, Suppl Table 4**). This is especially interesting because no non-endocrine markers were used in the deconvolution analysis. Instead, the non-endocrine proportion is the remainder signal left over after estimating the signal strength from each major endocrine cell type. Examination of the features most positively associated with non-endocrine proportion show highly biased expression in various non-endocrine cell types, across all omics layers. For example, GALK1 in proteomics (FDR = 1.6e-42; **Figure 4E, F**) is most expressed in acinar and ductal cells, *RBPMS* in RNA-seq (FDR = 0.00195; **Suppl Fig 3A**) is most expressed in ductal and stellate cells, and *MYC* in NanoString (FDR = 7.7e-6; **Suppl Fig 3B**) is most expressed in stellate cells.

**Figure 4.**
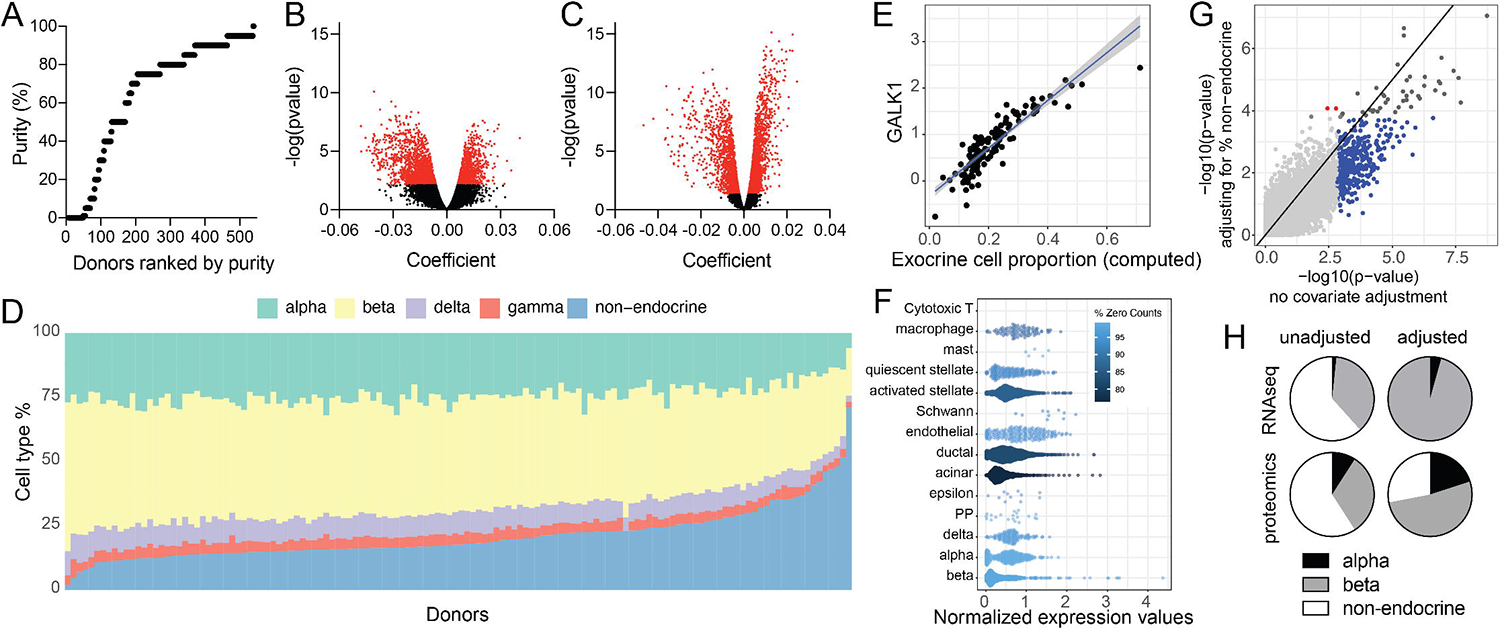
Effect of preparation purity on transcript and protein expression, and the use of cell type deconvolution analysis in correction for exocrine contamination. **A)** Estimate purity of preparations following isolation, with donors ranked in order by purity. **B-C)** Although islets are hand-picked prior to these analyses, post-isolation purity has an impact on both transcript (B) and protein (C) expression in human islets. Adjusted p-value <0.05 indicated in red. **D)** Cell type distributions computed from proteomic data across donors, ranked by ‘non-endocrine’ cell proportions. **E-F)** Correlation of GALK1 protein with the proportion of non-endocrine cells (E) and confirmation in single-cell RNA-seq data that *GALK1* is expressed primarily in acinar and ductal cells (F). **G)** Impact of adjusting for the proportion of non-endocrine cells on protein-level p-values for the association of bulk proteomics data and diabetes status (T2D vs. None). Proteins were either significant (FDR<0.05) both with and without adjusting for non-endocrine cells (dark grey), only when adjusting for non-endocrine cells (red), only when not adjusting for non-endocrine cells (blue), or not significant in either scenario (light grey). **H)** Of α-, β- and non-endocrine-associated RNA-seq and proteomics features found to be significantly different (adjusted p-value <0.05) when comparing T2D versus no diabetes with and without adjustment for the computed proportion of non-endocrine cell types (non-adjusted RNA-seq, n=117; adjusted RNA-seq, n=90; non-adjusted proteomics, n=134; adjusted proteomics, n=134).

**Table 2.** Features positively associated (FDR < 0.05) with each estimated cell type proportion. Beta, alpha, delta, and PP proportions are the % of all endocrine cells, and the association analysis for these cell types included non-endocrine % as a covariate. The percentage of features that were used as marker genes is shown in brackets.

| Cell type | Proteomics | RNA-seq | Nanostring |
| --- | --- | --- | --- |
| Non-endocrine % | 1491 (0%) | 849 (0%) | 20 (0%) |
| Beta cell % | 352 (5%) | 312 (11%) | 3 (0%) |
| Alpha cell % | 652 (10%) | 168 (13%) | 7 (57%) |
| Delta cell % | 1036 (1%) | 0 (0%) | 0 (0%) |
| PP cell % | 721 (1%) | 0 (0%) | 0 (0%) |

The α- and β-cell proportions are positively associated with many features in all three bulk omics datasets, most of which are not marker genes that were used in the deconvolution (**Table 2**, **Suppl Table 4**). Features with the most significant positive associations with the estimated α- and β-cell proportions have distinctive gene expression in the associated cell type, even for non-marker genes. For example, the non-marker gene features with the most significant positive associations with α-cell proportions are DOP1B (proteomics) and *LAT2* (bulk RNA-seq) and with β-cell proportions are SLC27A2 (proteomics) and *HHATL* (bulk RNA-seq).

Examination of the single cell distributions for these genes confirm the cell type-specific expression (**Suppl Figures 4 and 5**). Overall, there were more proteomics features associated with the estimated proportions than transcripts, and with lower p-values, even after excluding features corresponding to the marker genes used in the deconvolution analysis. This is likely because the proteomics data was used for the deconvolution, and therefore had the highest number of overlapping cell type proportion estimates and omics profiles (n = 134 for proteomics versus n = 90 and 91 for bulk RNA-seq and NanoString respectively), giving this analysis higher statistical power. Additionally, each omics dataset was from separate islet preparations. While the cell type composition should be similar across samples from the same donor, this likely contributes to the fewer significant associations observed for the bulk RNA-seq and NanoString data. Finally, while many features were significantly associated with the proportion of delta and PP cells in the proteomics data (**Table 2**), there were no significant associations with these cell types in either the bulk RNA-seq or NanoString gene expression data. However, examination of the single-cell gene expression distributions of the transcripts most significantly associated with the estimated delta and PP cell proportions (raw p-values < 0.05) shows that many of these do have distinct expression patterns in appropriate cell types. For example, in the bulk RNA-seq data, *F5*, *LRFN5*, and *SST* were the most associated with δ-cells and *FGB* and *CACNA2D3* were two of the four features most associated with PP cells (**Suppl Figures 6 and 7**).

**Figure 5.**
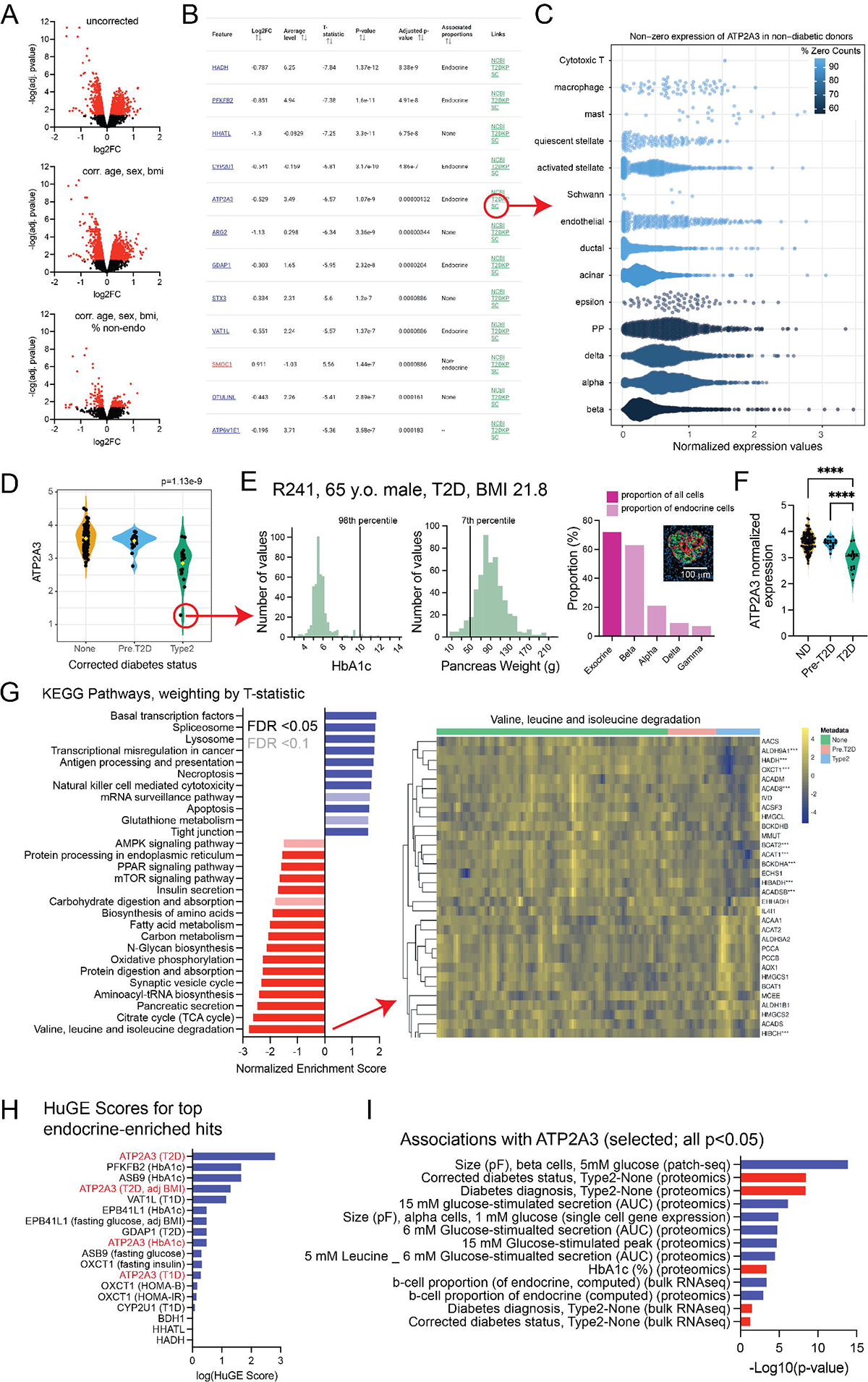
Mini-case study: Linking SERCA3b expression with diabetes and islet function. **A)** Analysis of proteomic data comparing the corrected diabetes status of ‘no diabetes’ versus ‘type 2 diabetes’. Analyses were run with no correction, with addition of age, sex and BMI as covariates, and then with addition of ‘proportion non-endocrine’ as a covariate. Adjusted p-value <0.05 indicated in red. **B)** Web-tool Results Table, with top hits sorted by adjusted p-value. **C)** Clicking on ‘SC’ (single-cell) raises a visualization of single-cell RNA-seq data for a particular hit (in this case *ATP2A3*) to confirm cell-type specific expression. **D)** Clicking on a row of the Results Table raises the protein expression of a hit (in this case ATP2A3) separated by the initial query. This example shows ATP2A3 expression in donors with no diabetes (ND), pre-type 2 diabetes (pre-T2D) and T2D (Type2). A potential outlier donor is indicated in the red circle. **E)** Clicking on the potential outlier raises the Donor View page for that donor for inspection. Shown here are the donor metadata, HbA1c and pancreas weight in relation to the total database donor population, and cell type composition for this potential outlier (R241). Composition calculations are from the proteomics deconvolution, with exocrine cells shown as a proportion of all cells and endocrine cell types shown as a proportion of all endocrine cells. Inset, the presence of both insulin (green) and glucagon (red) positive cells in R241 is confirmed by fluorescence immunostaining in banked FFPE biopsy from this donor. **F)** Proteomic data downloaded from the Data Download page, can be analyzed by the user. In this case the potential outlier was removed from analysis of ATP2A3 expression (****-p<0.0001; one-way ANOVA followed by Tukey post-test to compare groups). **G)** Gene set enrichment analysis, via the ‘pathway analysis’ tab highlights pathways up-regulated (blue) and down-regulated (red) in T2D. Results were exported in table format and plotted using local software. Within the tool, clicking on a pathway result raises a heatmap showing genes/proteins contributing to the pathway. Significant hits are indicated by ***. **H)** In the results table (panel B), clicking on ‘NCBI’ takes users to the NCBI gene page for a given hit, while clicking on ‘T2DKP’ takes users to the T2D Knowledge Portal page. Data extracted from the T2DKP page for top endocrine-enriched hits demonstrates Human Genetic Evidence (HuGE) scores suggesting potential links to human metabolism and diabetes. Red shows HuGE scores for *ATP2A3*. **I)** In the results table (panel B), clicking on the hit name (in this case ATP2A3) takes users to the Feature View results for that particular hit. Here, ATP2A3 is seen to be negatively (red) associated with T2D at both the transcript and protein levels, and positively (blue) associated with some measures of beta-cell function and insulin secretion.

### Control for purity in data analysis

Including the ‘non-endocrine percentage’ as a covariate greatly reduces the number of bulk omics features identified as statistically significantly different between donors with T2D and those with no diabetes, which is consistent with previous findings^37^ (**Figure 4G**). The significant feature count decreases from 1519 to 281 in the proteomics data, 418 to 43 in the RNA-seq data, and 25 to 10 in the NanoString data.

Importantly, adjusting for the non-endocrine proportion eliminates or greatly reduces statistically significant features with non-endocrine biased expression from the significant RNA-seq and proteomics results (**Figure 4H**), suggesting that this approach successfully filters out signals driven by differential amounts of trapped non-endocrine tissue. However, this correction also likely decreases the statistical power for detecting features that are truly reflective of perturbed islet function and could therefore be overly stringent.

### Mini Case-study: SERCA pump expression, genetic risk for diabetes, and islet function

To illustrate a potential analysis workflow that considers the different metadata factors, we first examined proteomic data comparing ‘Corrected diabetes status’ of ‘Type 2’ vs ‘No Diabetes’. As opposed to ‘Diabetes diagnosis’ which is based on diagnosis information provided upon organ procurement, the ‘Corrected diabetes status’ additionally accounts for donor HbA1c. Donors with reported T2D remain in that category, donors with no reported T2D, but HbA1c>6.5 are now considered as T2D, and donors with no diabetes and HbA1c between 5.7-6.5 are considered ‘pre-T2D’. With no covariate correction, this comparison yields 1636 differentially expressed proteins, while correcting for age, sex, and BMI yields 1478 (FDR<0.05). Because islets isolated from donors with T2D typically have lower purity as noted above,^38,39^ we further adjusted for the calculated ‘non-endocrine cell proportion’. This resulted in 284 significantly different proteins (138 up, 146 down; FDR<0.05; **Figure 5A**).

Many of the top hits, some of which were identified in the initial analysis of these data,^33^ appear enriched in endocrine cells (calculated from the proteomics deconvolution as above; **Figure 5B**). The results and analysis pipeline for this Feature View can be downloaded directly via the ‘Download Results’ and ‘Save Analysis’ buttons. This endocrine enrichment can be visualized at the single-cell RNA-seq level by clicking on ‘SC’ (shown for *ATP2A3*, **Figure 5C**), and information from the NCBI gene database for each hit can be accessed via the ‘NCBI’ link. For example, ATP2A3 (ATPase sarcoplasmic/endoplasmic reticulum Ca^2+^ transporting 3 protein) is also known as SERCA3b, a Ca^2+^-ATPase involved in clearing cytosolic Ca^2+^ and has been suggested to play a role in regulated insulin secretion.^40–42^

We visualized the downregulation of ATP2A3 protein expression in islets from donors with T2D (as opposed to no diabetes or pre-T2D) by clicking on the appropriate row in the ‘Results Table’ (**Figure 5D**). Although this protein is downregulated in T2D, a possible outlier with very low expression can be seen in the T2D group. Clicking on this outlier will open a new tab showing the ‘Donor view’ page for this donor, for inspection. This donor (R241) is a 65 year old male with T2D and elevated HbA1c (9.9%) but has a very small pancreas (7^th^ percentile; seen in the ‘Isolation Information’ tab), and the islet preparation has high exocrine content and very low purity (**Figure 5E**).The computed cell composition for this donor suggests a high proportion of β-cells, which we confirmed by immunostaining of banked paraffin embedded biopsy from this donor (**Figure 5E, inset**). To analyze ATP2A3 protein expression data with this potential outlier removed, we: *1)* From the ‘Data Download’ page filtered donors by availability of proteomics data; *2)* Selected ‘Donor information’ and ‘Proteomics’ as data to download in .xlsx format, and clicked ‘download’; 3) Analyzed the data by ‘Corrected diabetes status’ (‘diagnosis.computed’ in the downloaded file), using including outlier removal by ROUT (Q = 5%) which removed one outlier in each of the ‘Pre-T2D’ and ‘T2D’ groups, including R241. Analysis following removal of these outliers confirmed a significant reduction in ATP2A3 expression in the T2D group (**Figure 5F**).

Via the ‘Pathway Analysis’ tab we performed gene-set enrichment analysis (GSEA) on the results shown in **Figure 5A** using T-statistic for weighting with an FDR cutoff of 0.1 and the KEGG library. This revealed 54 significantly enriched pathways that appear either up- or down-regulated in T2D islets. Pathways include the citrate cycle (TCA cycle), synaptic vesicle cycle, insulin secretion, lysosome, and antigen processing/presentation. To narrow potentially overlapping pathways the GSEA can be re-run with collapse redundant sets checked, which filters out gene sets with gene lists and enrichment patterns that are statistically indistinguishable from other gene sets in the library. Re-running GSEA with this setting reduced the number of significant pathways to 36 (**Figure 5G**, selected pathways shown). The most significant of these (FDR=2.0×10^-7^) is ‘valine, leucine and isoleucine degradation’, as denoted by the dark shade of green in the ridge plot and the adjusted p-value in the ‘Results Table’ tab. The individual features in this pathway can be explored by clicking on the ridge plot to generate a heatmap of all proteins in this pathway (**Figure 5G**). As for the protein features, the results and analysis pipeline for this Pathway Analysis can be downloaded directly via the ‘Download Results’ and ‘Save Analysis’ buttons.

To explore the evidence from human genetics for a role of variation in genes in these pathways in metabolic control, the ‘T2DKP’ link associated with each hit can be clicked. This opens the relevant page in the Type 2 Diabetes Knowledge Portal to allow examination of common and rare variant associations in the gene and Human Genetic Evidence (HuGE) scores,^43^ tissue specific gene expression, and other genetic information as described recently.^31^ Several of the endocrine-enriched hits above appear to have genetic support for a role in metabolism, with HuGE scores ranging from ‘Moderate’ (>3) to ‘Compelling’ (>350) (**Figure 5H**). ATP2A3 for example presents “*compelling”* evidence that it is the gene mediating genetic associations at the locus with T2D (HuGE score 641.77), along with evidence for common variant associations with T2D adjusted for BMI, HbA1c, and T1D. This is in line with candidate genes studies of variants associated with T2D which have been replicated in larger genome-wide studies.^44^

To explore whether ATP2A3 is associated with other features, the feature name in the associated results table can be clicked, or, after navigating to the ‘Feature View’ page, ATP2A3 can be directly searched. This reveals statistically significant associations of potential interest at both the protein and transcript levels, including a negative association with HbA1c (adjusted p-value = 4.4×10^-5^); down-regulation of *ATP2A3* transcript in T2D (adjusted p-value = 0.036); a positive association with β-cell size in Patch-seq data (adjusted p-value = 4.3×10^-12^); and positive associations with various measures of insulin secretion, such as the area under the curve for insulin secretion at 15 mM glucose (adjusted p-value = 7.6×10^-7^) (**Figure 5I**).

## DISCUSSION

The past decade has produced major efforts to generate large amounts and diverse types of data from human pancreas that can be leveraged for insights into diabetes pathophysiology and potential therapeutic targets. Access to, and integration of, these large datasets remain a challenge. Three main approaches have emerged to facilitate integrated human pancreas and islet datasets: 1) Portals that aggregate minimally processed datasets and images;^7,18,20,21,45,46^ 2) Provision of islet quality control and donor demographic data, often associated with biobanking activity;^19,47^ and 3) Implementation of pipelines for data analysis and visualization, largely in the genomic space.^22,25,48,49^ Each of these approaches are tremendously valuable in understanding islet and pancreas (patho)physiology, but naturally are subject to key limitations. Current resources occasionally include low numbers of donors, little or no donor or technical data, limited accessibility to underlying data, narrow breadth of data types, or conversely broad and deep datasets that require advanced expertise for analysis. The ADI IsletCore (www.isletcore.ca) is a large single-center biobanking program that has provided human islets for research since 2011.^39^ We have, over more than a decade, increased the amount of data collected from each research islet isolation through a collaborative ‘distributed phenotyping network’,^14,33,39,50,51^ and have now leveraged this to develop <u>HumanIslets.com</u> as an open resource for the community.

Our goal is to provide a flexible, open, and user-friendly platform for exploring the relationship between donor, isolation, and islet phenotypes and molecular datasets from many donors. This should be useful to the broader diabetes and islet research community via built-in analysis pipelines with standardized options and outputs, and for users who are more experienced in data processing and analysis by providing ease of access to minimally processed data in a variety of formats when possible. To balance data access, research benefits, and requirements for privacy and compliance, all data are de-identified at the source of organ procurement and sensitive data such as RNA sequencing, is provided only in a processed form while raw data remains accessible via controlled access repositories. Although the depth of pancreas phenotyping here does not presently match that provided by detailed case-control initiatives, notably the Human Pancreas Analysis Program (HPAP),^7,18^ our goal is to provide access to as broad a range of donors as possible, facilitating an understanding of both biological variation and diabetes pathophysiology. We also provide more detailed metadata describing not only donor phenotypes, but also organ processing parameters and cell culture outcomes which are not commonly included with publicly available datasets elsewhere.

Confounding relationships can be quickly viewed with other variables and across different omics modalities to inform hypotheses about causality. Visual analytics can help researchers assess potential relationships beyond the summary statistics. The technical parameters, especially, should be commonly available since they add no identifiable information. Indeed, our exploration of potential confounding technical variables led us to include culture time as covariate when generating precomputed results for the Feature View page. We did not include purity as a covariate since it is only a rough estimate of the amount of non-endocrine tissue in the analyzed samples, particularly following hand-picking, and often confounds measures of islet function. We also did not include the computed non-endocrine proportion, mainly because the estimates are only available for donors with proteomics data.

This can nonetheless be informative, and we include a column in the results table that indicates whether each feature is positively associated (FDR < 0.05) with either the estimated endocrine or non-endocrine proportions, to alert researchers to cases where a significant relationship could be explained by variable cell type composition. Also, the cell type proportions are available under the covariates drop-down menu in the omics Analysis page, so researchers can employ it in their own customized analyses if they choose.

This resource can replicate well-established findings from us and others. We highlighted commonly accepted pathophysiology above in **Figure 5**, for example reduced oxidative phosphorylation in T2D islets has previously been shown both at the transcript and functional levels.^52,53^ Analysis of single-α-cell patch-seq data via the web tool also replicates our previous findings^54^ that the expression of lineage marker transcripts (e.g. *ISL1*, *NKX2-2*, *NEUROD1*, *RFX6*) all negatively correlate with α-cell function in T2D, and with numerous developmental pathways (e.g. GSEA for Go Biological Process ‘Central Nervous System Development’, FDR=6.0×10^-4^; n=620 cells from 24 donors). It is important to note that some results obtained with this platform may not match exactly published findings using the same or overlapping datasets, likely due to the updated datasets and/or slightly different analysis pipelines.

In summary, <u>HumanIslets.com</u> is a valuable resource for the diabetes research community, containing data and utility both for wet lab scientists, with easy analysis and visualization of data at the donor-islet-cellular level, and for computational biologists, providing straightforward downloading of minimally processed datasets. This resource continues to evolve through the ongoing collection of datasets outlined in this report, and by the addition of new data types for additional analyses. Examples of the latter, include donor genotyping; genetically predicted ancestry, and genetic risk scores for diabetes;^55,56^ prohormone processing;^57,58^ metabolomics;^59^ and the pancreatic accumulation of environmental contaminants^60,61^ and lipids,^62^ among other data types that will be added in the future. In time, we hope to allow all ADI IsletCore islet and tissue recipients to provide/upload data for integration. We welcome feedback for future refinement of this resource (see ‘Feedback’ under the ‘About’ tab - https://www.humanislets.com/#/about), which we hope will support exploration and hypothesis generation by the diabetes, transplant, and metabolism research community.

### Limitations

We attempted to balance sophisticated *versus* intuitive analysis pipelines, to serve both inexperienced and experienced users and allow meaningful analysis within the computing restrictions of a web server. Some detailed analyses will require off-line computation, and while the data processing pipeline was designed for maximum comparability across omics types there may be more optimal analyses for each individual dataset. This should be facilitated, however, through the addition of a donor/data filtering and download tool. While the ability to easily add new and expanded datasets is an important advantage of this platform, it is important to note that some results will change as new data are added (i.e. pre-computed statistics will be different after they are re-computed with more samples). As such, version tracking and reporting by users will be important. While some datasets are limited by low total donor numbers or lack some overlap between measures (i.e. there is currently little overlap between oxygen consumption measurements and RNA-seq or proteomics results, as these remain in analyses pipelines), the continuing addition of data when available will enhance the overall functionality of the resource. Additionally, while this dataset aims to include as much technical and donor metadata as possible, some donor metadata that may be important are not available. This may include information on donor gender or reproductive status, ethnicity, and exposure history, among potentially many others. Some of our ongoing new dataset collection explicitly designed to fill these gaps include plasma measurement of sex hormones, determination of genetic ancestry, and pancreas accumulation of environmental contaminants. Finally, long-term maintenance of academic software and databases are challenging. However, our team has extensive experience in tool maintenance^63,64^ and tissue biobanking sustainability that should benefit the long-term evolution of <u>HumanIslets.com</u> in support of diabetes and islet research.

### Consortium

Members of the HumanIslets Data Analysis and Sharing (HI-DAS) Consortium are Ella Atlas (Health Canada), Austin Bautista (Alberta), Jennifer E Bruin (Carleton), Alice L Carr (Alberta), Haoning H Cen (UBC), Yi-chun Chen (UBC), Angela Ching (Carleton), Xiao-Qing Dai (Alberta), Tina Dafoe (Alberta), Theodore dos Santos (Alberta), Cara E Ellis (Alberta), Jessica D Ewald (Broad Institute), Leonard J Foster (UBC), Anna L Gloyn (Stanford), Aryana Hossein (UBC), Myriam P Hoyeck (Carleton), James D Johnson (UBC), Jelena Kolic (UBC), Yao Lu (McGill), Francis Lynn (UBC), James G Lyon (Alberta), Patrick E MacDonald (Alberta), Jocelyn E Manning Fox (Alberta), Renata Moravcova (UBC), Nadya M Morrow (Ottawa), Erin E Mulvihill (Ottawa), Andrew Pepper (Alberta), Lindsay Pallo (UBC), Varsha Rajesh (Stanford), Jason Rogalski (UBC), Shugo Sasaki (UBC), Seth A Sharp (Stanford), Nancy P Smith (Alberta), Aliya F Spigelman (Alberta), Han Sun (Stanford), Swaraj Thaman (Stanford), C Bruce Verchere (UBC), Jane Velghe (UBC), Charles Viau (McGill), Kyle van Allen (Carleton), Jessica Worton (Alberta), Jordan Wong (Alberta), Jianguo Xia (McGill), Dahai Zhang (UBC).

## Methods

A general overview of methods and usage of the HumanIslets.com web tool can be found in the online documentation (http://doc.humanislets.com) and as **Supplementary Document 2**. Subsets of all data types have been published elsewhere, with extensive details on the protocols used to perform experiments and collect the data. Here, we provide a brief overview of the entire process, along with references to the previous descriptions of complete data collection methodology.

### Islet isolation and culture

Islets were isolated from pancreas of cadaveric human donors at the Alberta Diabetes Institute IsletCore using standard procedures.^39,65,66^ The isolated islets were cultured prior to experimental use or distribution in CMRL 1066 (Corning) supplemented with 0.5% BSA (Equitech-Bio), 1% insulin-transferrin-selenium (Corning), 100 U/mL penicillin/streptomycin (Life Technologies), and L-glutamine (Sigma-Aldrich) at 22°C with 5% CO_2_ at a typical seeding density of 225 IEQ/cm^2^. Following release from the ADI IsletCore, islets were typically cultured in DMEM (11885, Gibco) supplemented with L-glutamine, 110 mg/l sodium pyruvate, 10% fetal bovine serum (FBS) (12483, Gibco), and 100 U/ml penicillin/streptomycin (15140, Gibco) at 37°C with 5% CO_2_.

All research was performed with approval of the Human Research Ethics Board at the University of Alberta (Pro00013094; Pro00001754), the University of British Columbia (UBC) Clinical Research Ethics Board (H13-01865), the University of Oxford’s Oxford Tropical Research Ethics Committee (OxTREC Ref.2-15), the Oxfordshire Regional Ethics Committee B (REC reference: 09/H0605/2), or by the Stanford Center for Biomedical Ethics (IRB Protocol: 57310). All donors’ families gave informed consent for the use of pancreatic tissue in research.

### Insulin secretion

Static glucose-stimulated insulin secretion was performed as described.^67^ Briefly, on the day of sample collection islets were hand-picked and cultured overnight. Fifteen hand-picked islets, in triplicate, from each donor were pre-incubated at low glucose for two 1-hour periods, consecutively, and then sequentially treated with low glucose followed by high glucose 1 hour each. Experiments were performed with parings of glucose concentrations as follows: 1→10mM; 1→16.7mM; and 2.8→16.7 mM. Collected supernatants, and insulin content collected by acid ethanol lysis of the pellet, were measured by ELISA (ALPCO STELLUX Chemiluminescence). For in-tool omics Analysis, a single donor-level measurement was computed as the median of the three replicate measures, reducing the impact of replicate-level outliers. Then, donor-level values were filtered to remove those that were more than 4 standard deviations away from the mean of all donors on a log10 scale. This removed values that were many orders of magnitude different from the population. Data in the Data Download tool are provided at the replicate level (before outlier removal). Dynamic insulin secretion responses were assessed as described^68^ following shipment of islets in CMRL media (Thermo Fisher Scientific) overnight from Edmonton to Vancouver.

Islets were hand-picked and cultured in RPMI 1640 medium (Thermo Fisher Scientific, Cat#, 11879-020) supplemented with 5.5 mmol glucose, 10% FBS, and 100 units/mL penicillin/streptomycin for 24-72 hours prior to experiments to allow for recovery from shipment. Perifusion columns were loaded with 65 islets and perifused at 0.4 mL/min with 3 mM glucose Krebs-Ringer Modified Buffer (KRB) solution for 1-hour. Islets were perifused with 6 mM or 15 mM glucose, and subsequently 30 mM KCl. Parallel experiments examined responses to 5 mM leucine (Sigma, Cat. # L8912, dissolved in 1M HCl, then pH adjusted with 1M NaOH) and a 1:1 mixture of 0.75 mM oleic acid (Sigma, Cat. # 364525) and 0.75 mM palmitic acid (Sigma, Cat. #P5585) at either 3 mM or 6 mM glucose (see figure legends). Fatty acids stock solutions were prepared in 50% ethanol at 65°C for 30 minutes, and then added to a solution of fatty acid free bovine serum albumin (BSA) (Sigma, Cat. # A7030) in a 6:1 molar ratio in a 37 °C water bath for 60 min. Peak insulin secretion was defined as single point whereby the amount of insulin released during the first 15 minutes of a solution change was the highest. Samples were stored at –20 °C and insulin secretion was quantified using human (Millipore Cat. # HI-14K) insulin radioimmunoassay kits.

### Oxygen consumption

Agilent’s Seahorse XFE24 Analyzer with the islet capture microplates was used for oxygen consumption measurements. On the day of sample collection, islets were hand-picked to 70 islets per well in triplicate. Islets were placed in the center of the well, with islet capture screens used to keep islets in place, and incubated at 37°C for 1 hour without CO_2_. Islets were exposed to a modified Mito Stress Test (Seahorse XF Cell Mito Stress Test Kit, Cat. No. 103015-100) where basal measurement was at 2.8 mM glucose and stimulation at 16.7 mM glucose, then with 5 µM oligomycin, 3 µM FCCP, and 5 µM of rotenone and antimycin A. All solutions were made using DMEM (from Agilent) supplemented with 1% FBS, sodium pyruvate, and L-glutamine. After the run was complete, islets were collected and a DNA assay (QuantiFluor® dsDNA System, Promega, Madison, WI, USA) was conducted for data normalization.

### Electrophysiology

was performed as described.^54,69^ Islets were dispersed to single cells, plated as single cells in 35-mm dishes, and cultured for 1-3 days in RPMI with 5 mM glucose (37°C, 5% CO_2_). Patch-clamp was in the whole-cell configuration using a HEKA EPC-9 amplifier (Harvard Apparatus) with 1, 5 or 10 mM glucose. Cells were held at −70 mV and depolarizations were applied to cells to elicit activation of voltage-dependent Na^+^ and Ca^2+^ channels, and exocytosis responses were measured as increases in whole-cell capacitance. Responses are normalized to initial cell size. Following electrophysiological measurements, cells were identified by post-hoc immunostaining for insulin with a rabbit anti-insulin primary antibody (Santa Cruz; #SC-9168; RRID: AB_2126540) and goat anti-rabbit Alexa Fluor488 secondary (ThermoFisher, #A-11076; RRID: AB_141930), and with a guinea pig anti-glucagon primary antibody (Sigma-Aldrich, #G2654; RRID: AB_259852) and goat anti-guinea pig Alexa Fluor 594 secondary (ThermoFisher, #A-11076; RRID: AB_141930); or following collection for scRNA-seq.

### Omics data processing

#### Bulk RNA-seq

150 human islets were hand-picked on the day of distribution and placed in 1ml of Trizol (Invitrogen), vortexed and stored at −80°C until shipping to Oxford or Stanford for further processing. Ensembl gene IDs were mapped to Entrez IDs and counts from any duplicated Entrez IDs were summed. Transcripts with over 80% zeros were filtered out. The RNA-seq dataset contains seven batches, and significant batch effects were observed. Batch effects were corrected for using the Combat-Seq function in the *sva* R package (version 3.44.0)^70^, which preserves the count nature of the data. Counts were converted to log-counts-per-million (logCPM) using the relative log expression (RLE) method in the *edgeR* R package (version 3.38.4).^71^ Transcripts with logCPM variance in the lowest 20th percentile were filtered out. Since the sequencing depth was different across different batches, a substantial number of transcripts had zero counts in some batches and low but robust counts in other batches. These batch-related drop-out patterns led to large numbers of significant differentially expressed features that disappeared if individual batches were held out. To address this problem, counts with a value of zero or one were replaced with NAs, excluding these individual values from all downstream statistical analyses and leading to more stable differential expression analysis results.

#### PatchSeq

Reads from FASTQ files (available in the NCBI Gene Expression Omnibus (GEO) and Sequence Read Archive (SRA) under accession numbers GSE124742,^69^ GSE164875,^54^ at PancDB,^72^ and unpublished data to be deposited) were trimmed using TrimGalore version 0.6.5 with auto-detection of adapter sequence, then aligned and counted using STAR version 2.7.9a to the GRCh38 human genome (Ensembl release 104). Cell count matrices were loaded into R^73^ and a Seurat object was generated using the *Seurat* R package (version 4.3.0).^74^ Only cells with patch-clamp data were included; data for additional cells are available from the listed accession numbers. Cell types were identified by projection, using the MapQuery command from the R package *Seurat*, onto a large set of human islet single cell RNA-seq data generated from publicly available data (see below). Cells were filtered to remove any that were not α-or β-cells, had fewer than 500 total counts across all transcripts, had fewer than 300 unique transcripts with non-zero count values, or had greater than 20% mitochondrial DNA. Ensembl gene IDs were mapped to Entrez IDs and counts from any duplicated Entrez IDs were summed. Counts were separated by cell type (α- and β-) and glucose concentration (1, 5, and 10mM), resulting in six different matrices. Each matrix was filtered to remove transcripts with greater than 80% zeros. Counts were converted to logCPM values using the trimmed mean of M-values (TMM) method from the *edgeR* R package (version 3.38.4).^71^ TMM has been shown to perform better for single-cell RNA-seq data compared to the RLE method because it is robust to the high drop-out rate. Individual transcripts that had zero counts were replaced with NA. Electrophysiology parameters other than the half-inactivation sodium current were normalized to cell size. The sign of the calcium entry, sodium current, and calcium current (early and late) electrophysiology parameters was flipped such that greater values correspond to greater currents. Negative exocytosis values were replaced with zeros.

#### Pseudobulk RNA-seq

The single-cell counts described above were also summarized at the donor level using a pseudobulk approach. Counts were separated by cell type (α- and β-), creating two matrices. For each matrix, the number of cells per donor was counted, and only cells from donors with at least five cells were retained. Pseudobulk profiles were created by summing counts from all cells derived from the same donor. Ensembl gene IDs were mapped to Entrez IDs and counts from any duplicated Entrez IDs were summed. Each matrix was filtered to remove transcripts with greater than 80% zeros. Counts were converted to logCPM values using the TMM method from the *edgeR* R package (version 3.38.4).^71^ Individual transcripts that had zero or one count were replaced with NA values.

#### NanoString

Fifty islets were hand-picked on the day of distribution and placed in 100 µl RLT (Qiagen) with 1% beta mercaptoethanol (Sigma) and stored at −80°C until shipping to the University of British Columbia for processing. Briefly, 1.5 µl cell lysate was used for hybridization with customized gene code sets for 16 hours. The hybridized complex was then loaded on one cartridge, and the NanoString nCounter Sprint Profiler was used to measure the target gene expression. Data are analyzed with nSolver4.0 software for quality control and normalization to a panel of housekeeping genes (NanoString, USA). NanoString data were collected from two different gene lists, the first with 132 transcripts and the second with 147 transcripts, and 156 unique transcripts across both. Significant batch effects were present, and so transcripts measured in both NanoString arrays were batch corrected using the Combat function from the *sva* R package (version 3.44.0).^70^ Given the much smaller number of transcripts compared to the RNA-seq data, no abundance or variance filters were applied.

#### Bulk proteomics

Islets were shipped to the University of British Columbia, hand-picked, and cultured in RPMI for 24-72 hours. After recovery, 300 islets were hand-picked again, washed with PBS, snap frozen as a pellet, and stored at −80°C. LC-MS/MS analysis was performed with n = 3 technical replicates, using a NanoElute UHPLC system (Bruker Daltonics) with Aurora Series Gen2 (CSI) analytical column coupled to a Trapped Ion Mobility-Time of Flight mass spectrometer (timsTOF Pro; Bruker Daltonics, Germany) operated in Data-Independent Acquisition-Parallel Accumulation-Serial Fragmentation (DIA-PASEF) mode. Protein abundances were filtered to remove any that had more than 50% missing values, median normalized, and log2 transformed. Missing values were imputed with the missForest method (random forest-based) using the *imp4p* R package (version 1.2.0).^75^ Uniprot IDs were converted to Entrez IDs, and rows with duplicate IDs were aggregated by taking the mean. Proteins with variance in the lowest 20th percentile were filtered out.

For all omics datasets, both the minimally and fully processed matrices were saved so that they can be accessed by researchers on the Data Download page. Minimally processed datasets are the data in matrix form, but with no filters, normalization, missing value imputation, or batch correction applied. Raw data such as FASTQ files for the RNA-seq are not available for download through the tool. Instead, accession IDs indicating where they are available on public repositories are available in the donor metadata file (“RRID” column), or at the accessions listed in the data accessibility section below.

### Statistical analysis

The following methods are called by the omics Analysis page to identify omics features significantly associated with selected metadata and pathways overrepresented or enriched in the association results.

#### Feature association analysis

Linear models are used to compute omics feature-metadata associations for the bulk RNA-seq, NanoString, pseudobulk RNA-seq, and bulk proteomics datasets. A linear model containing the selected metadata variable of interest and all covariates (if any) is fit to the expression or abundance levels of each feature in the selected omics dataset, using the *limma* R package (version 3.52.4).^76^ If the user selects a categorical metadata of interest, two dropdowns automatically appear so that a specific contrast can be specified (ie. type 2 diabetes versus no diabetes). Then, the coefficient of the metadata variable (continuous) or metadata contrast (categorical) of interest and its associated p-value are extracted from the model for each feature. P-values are adjusted using the false discovery rate method. The rationale for using the original *limma* method is that it has been shown to perform well for many omics types as long as the data are appropriately normalized and transformed.

While more customized methods developed for specific omics types may perform slightly better when applied to the appropriate data type, the advantage of our approach is that it can be applied to many omics types without modification, making the results more comparable to each other and more consistent for users to interpret.

Due to the highly nonlinear nature of the relationships between single-cell gene expression and single-cell electrophysiology outcomes, Spearman correlation is used to compute feature-metadata relationships for the patchSeq data. Only single-cell electrophysiology outcomes are available for analysis with the single-cell expression data, and covariate adjustment is not supported. The single-cell expression profiles are also summarized as pseudobulk profiles for each donor; associations with donor-level clinical, technical, and functional outcomes can be investigated using this format.

#### Pathway analysis

Overrepresentation analysis (ORA) and gene set enrichment analysis (GSEA) are supported for pathway analysis of the feature association results. Both methods are implemented using the *fgsea* R package (version 1.22.0).^77^ The feature list used for ORA is determined by the p-value threshold specified in the association analysis. GSEA can be performed using either the model coefficient (equivalent to log2FC when the metadata is categorical) or the test statistic from the association analysis to rank the features. For the patchSeq data, the coefficient is the correlation estimate and there is no test statistic option as it is not compatible with the Spearman correlation results. There are six supported pathway libraries, including KEGG, Reactome, the full Gene Ontology term list (broken into Biological Process, Cellular Component, and Molecular Function), and the PANTHER summary of the Gene Ontology terms (also broken into the same three categories). Pathway analysis can sometimes return a high number of significant pathways or gene sets that consist of mostly the same features, leading to a redundant results list. Users have the option of collapsing redundant sets, which uses the “collapsePathways” function within the *fgsea* R package to filter the list of gene sets to only include those that have statistically significantly distinct feature lists. Pathway analysis is not supported for the NanoString data, as there are too few features (*n* = 156) for adequate pathway coverage.

#### Pre-computed Results

Associations were computed between all omics feature-metadata variable pairs and between all metadata variable-metadata variable pairs. The linear model and Spearman correlation methods described above were used to compute the omics-metadata associations, with one modification. Some categorical variables have obvious specific contrasts, for example for diabetes status, one is usually interested in type 1 versus no diabetes and type 2 versus no diabetes. In these cases, we specify each contrast of interest using the methods described above. Other variables do not have obvious comparisons, for example donation type, so in this case an ANOVA style analysis is done and the statistics are extracted for the categorical variable coefficient rather than for a specific contrast. Sex, age, BMI, and culture time were included as covariates for analyses that used a linear model (all bulk omics data associations).

Three different methods were used to compute associations between pairs of metadata, depending on whether each metadata in the pair were continuous or categorical. Pearson correlation was used for continuous-continuous pairs, Cramer’s V correlation was used for categorical-categorical pairs, and ANOVA was used for categorical-continuous pairs where the categorical variable is the independent variable, and the continuous variable is the dependent variable. Multiple hypothesis correction (FDR method) was performed across all results for the same statistical method. Some metadata variables such as percent purity and percent trapped are in between ordinal variables and a true continuous variable because they are estimates made by eye by a research technician, and the technician tends to estimate whole percentages (ie. 80%, 70%). For simplicity, we treat these as continuous variables.

#### Single-cell database compilation

Reads from FASTQ files (available in the NCBI Gene Expression Omnibus (GEO) and Sequence Read Archive (SRA) under accession numbers GSE81076,^78^ GSE81547,^79^ GSE81608,^80^ GSE83139,^81^ GSE84133,^37^ GSE85241,^82^ GSE86469,^83^ GSE154126;^84^ from the European Molecular Biology Laboratory European Biology Institute (EMBL-EBI) under accession number E-MTAB-5061;^85^ at PancDB up to donor HPAP-109^72^) were trimmed using TrimGalore version 0.6.5 with auto-detection of adapter sequence, then aligned to the GRCh38 human genome, Ensembl release 104,and counted using STAR version 2.7.9a except for files from GSE84133, which were aligned and counted using the inDrops protocol; files from GSE81076 and GSE85241, which were aligned and counted using the R package *scruff* (version 1.10.1)^86^ with *Rsubread* (version 1.22.2);^87^ and files to be deposited (unpublished), which were aligned and counted with Cell Ranger (10X Genomics Cell Ranger v6.1.2). Cell count matrices were loaded into R^73^ and objects generated using the R package *Seurat (*version 4.3.0),^74^ restricted to transcripts detected in more than 10 cells and cells with between 200 and 12,500 unique transcripts with non-zero count values; cells with percent mitochondrial transcripts in the top 25% for each dataset were also dropped. Transcript names were maintained as Ensembl identifiers to preserve the greatest number of unique transcripts. After combining all donors, transcripts expressed in fewer than 0.1% of cells were filtered. Donor HPAP-027 was dropped from the data set because there were very few cells (32 cells). The reciprocal PCA intergration with references (HPAP-037, HPAP-040; HPAP-045, HPAP-050, HPAP-052, and HPAP-072) workflow from the R package *Seurat* (version 4.3.0)^74^ was followed, with k.anchor = 5, then FindNeighbors and FindCluster (resolution = 1.2). Cell types were first manually annotated based the unambiguous, high level expression of marker genes (**Suppl Table 5**) in distinct clusters in UMAP space, except cytotoxic T cells, which did not form a distinct cluster but had restricted expression of *TRAC*. The remaining ambiguous cells were annotated using the MapQuery function by projecting unknown cells onto the manually annotated cells.

#### Deconvolution analysis

After normalization for tissue sample size, the intensity of a marker protein that is only abundant in one cell type and that has relatively constant levels is proportional to the abundance of that cell type. High-quality lists of marker genes have been published for islet endocrine cell types based on a meta-analysis of seven different human islet single-cell datasets.^88^ Studies have also published the range of endocrine cell type proportions across human donors, as assessed by staining and microscopy - here, we assume an average of 0.55 β-cells, 0.30 α-cells, 0.10 8-cells, and 0.05 PP cells.^89,90^ We used this prior knowledge to estimate the cell type composition of each donor using the proteomics data. First, islet endocrine marker gene lists from van Gurp et al.^88^ were filtered to remove any that were classified as markers for more than one cell type and any that were not present in the proteomics data. This resulted in 72 genes for β-cells, 78 for α-cells, 15 for δ-cells, and 8 for PP cells. Next, we computed the proportion of each cell type for each donor, according to each marker gene (*prob*_*c,m*_). This was done using the following formula:

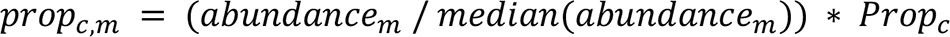

For example, the proportion of β-cells in a given donor according to the marker gene *INS* would be computed as the abundance of insulin, divided by the median insulin abundance across all donors, and multiplied by the prior known median proportion of β-cells (0.55). The overall proportion of each cell type was calculated as the median of proportion estimates across all marker genes for that cell type. Finally, the proportion of non-endocrine cells was computed as the remainder after adding the estimated proportion of β-, α-, δ-, and PP cells and normalizing the endocrine proportions to satisfy an assumed maximum purity of 98%. When computing associations between individual omics features and cell type proportions, endocrine cell proportions were normalized as a proportion of the total endocrine signal and the non-endocrine proportion was included as a covariate to better distinguish cell type-specific signals.

#### Tool Implementation

The *HumanIslets* frontend is implemented using Angular version 14, with the PrimeNg components library. All statistical analyses are performed in R, are managed by a Java backend with Rserve, and are communicated from the frontend to the backend via REST API. Interactive visualizations are implemented with d3 in the frontend and static visualizations are generated in the backend using the *ggplot* R package (version 3.4.2).^91^ The R functions used to generate all statistical results and visualizations are available at the HumanIsletsR Github repository (https://github.com/xia-lab/HumanIsletsR).

## Data accessibility

All raw data used in this study is available for download via the *HumanIslets* web-tool, with the following exceptions for raw sequencing and proteomics data (available elsewhere). Bulk RNA sequencing data and raw proteomics data are from a recent preprint^33^ and has been deposited in the European Genome-phenome Archive (EGA; <u>EGAS00001007241</u>) and ProteomeXchange via MASSive (PXD045422), respectively. Patch-seq sequencing reads are from published papers,^54,69^ new data, and the Human Pancreas Analysis Program (HPAP) and is available in the NCBI Gene Expression Omnibus (GEO) and Sequence Read Archive (SRA) under accession numbers <u>GSE124742</u>, <u>GSE164875</u>, and at the PANC-DB data portal of the HPAP (https://hpap.pmacs.upenn.edu),^72^ along with data to be deposited (unpublished). Data used to generate islet cell type-specific single-cell expression profiles is from HPAP (availability as above) and, from new data (GSE269204), and from published papers.^37,78–85^ The latter are available via NCBI GEO and SRA via accession numbers GSE81076, GSE81547, GSE81608, GSE83139, GSE84133, GSE85241, GSE86469, GSE154126, and from the European Molecular Biology Laboratory European Biology Institute (EMBL-EBI) under accession number E-MTAB-5061.

## ACKNOWLEDGEMENTS

The University of Alberta acknowledges that we are located on Treaty 6 territory, and respects the histories, languages, and cultures of First Nations, Métis, Inuit, and all First Peoples of Canada, whose presence continues to enrich our vibrant community. We thank all families and donors for the generous gifts in support of diabetes and transplantation research, and the Human Organ Procurement and Exchange (HOPE) program, Southern Alberta Organ and Tissue Donation Program (SAOTDP), Trillium Gift of Life Network (TGLN), BC Transplant, Quebec Transplant, and other Canadian organ procurement organizations for their efforts procuring pancreas for research. We appreciate the input of Viljem Pohorec (University of Maribor), Joan Camunas-Soler (Gothenberg), Jon Campbell (Duke University) and Jane Velghe (UBC).

