## Supplemental Figures for "HumanIslets: An integrated platform for human islet data access and analysis"

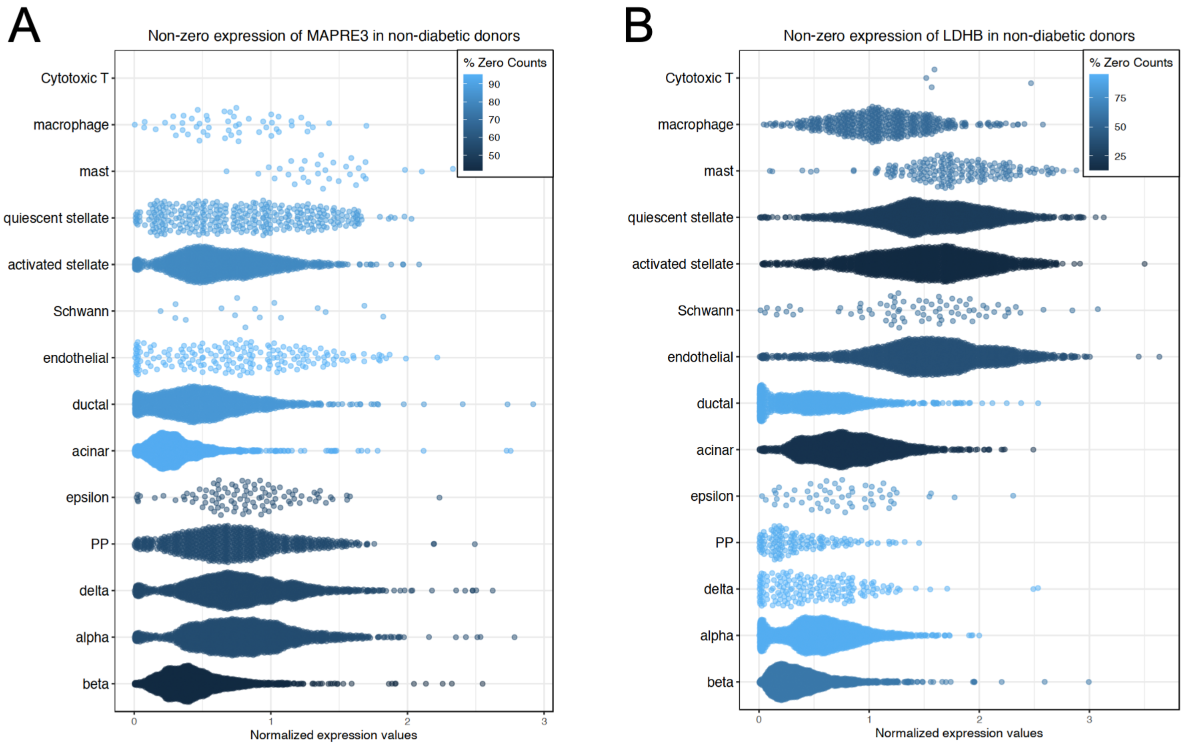


**Suppl Figure 1.** Bulk proteomics features with the most significant positive (**A**; MAPRE3, coef = 0.013, FDR = 3.4e-12) and negative (**B**; LDHB, coef = -0.015, FDR = 2.4e-10) associations with the islet % purity.

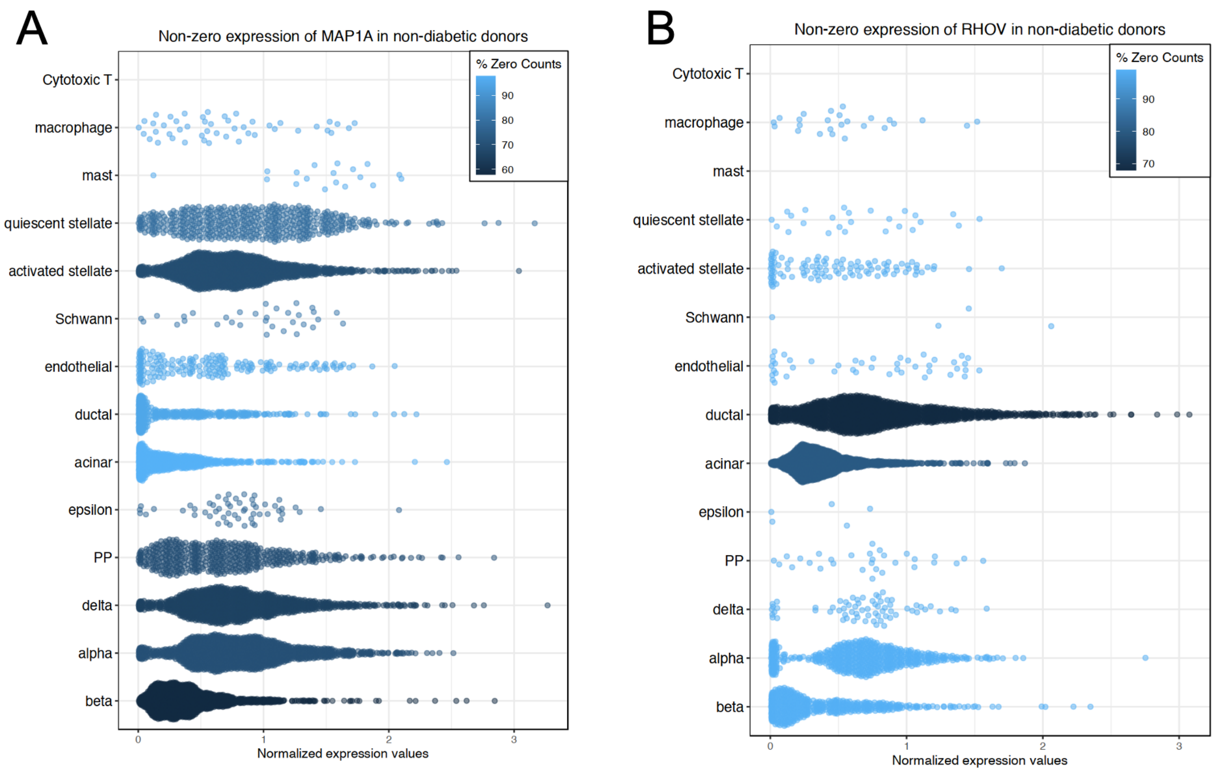


**Suppl Figure 2.** Bulk RNA-seq features with the most significant positive (**A**; MAP1A, coef = 0.020, FDR = 2.3e-5) and negative (**B**; RHOV, coef = -0.041, FDR = 1.2e-6) associations with the islet % purity.

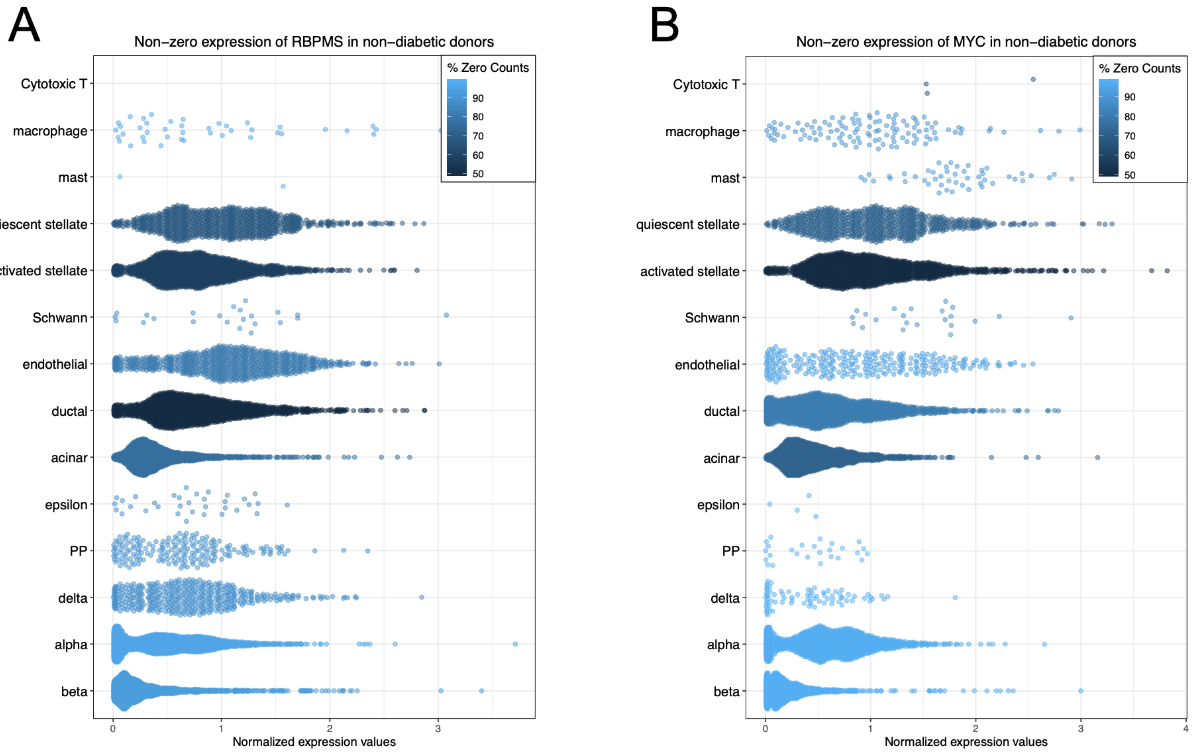


**Suppl Figure 3.** Bulk RNA-seq (**A**; RBPMS, coef = 2.70, FDR = 0.0020) and Nanostring (**B**; MYC, coef = 4.2, FDR = 7.7e-6) with the most significant positive associations with non-endocrine proportion.

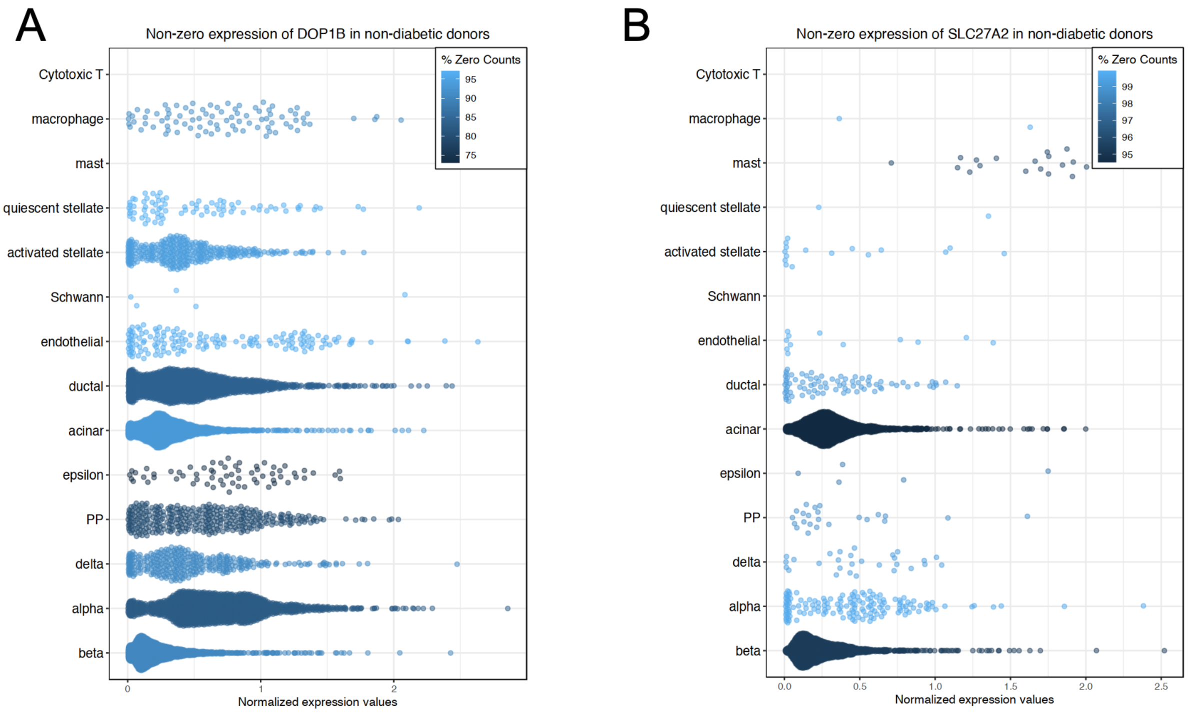


**Suppl Figure 4.** Proteomics features with the most significant positive associations with alpha cells (**A**; DOP1B, coef = 6.71, FDR = 1.7e-16) and beta cells (**B**; SLC27A2, coef = 5.14, FDR = 3.3e-11) that were not included as marker genes in the deconvolution analysis.

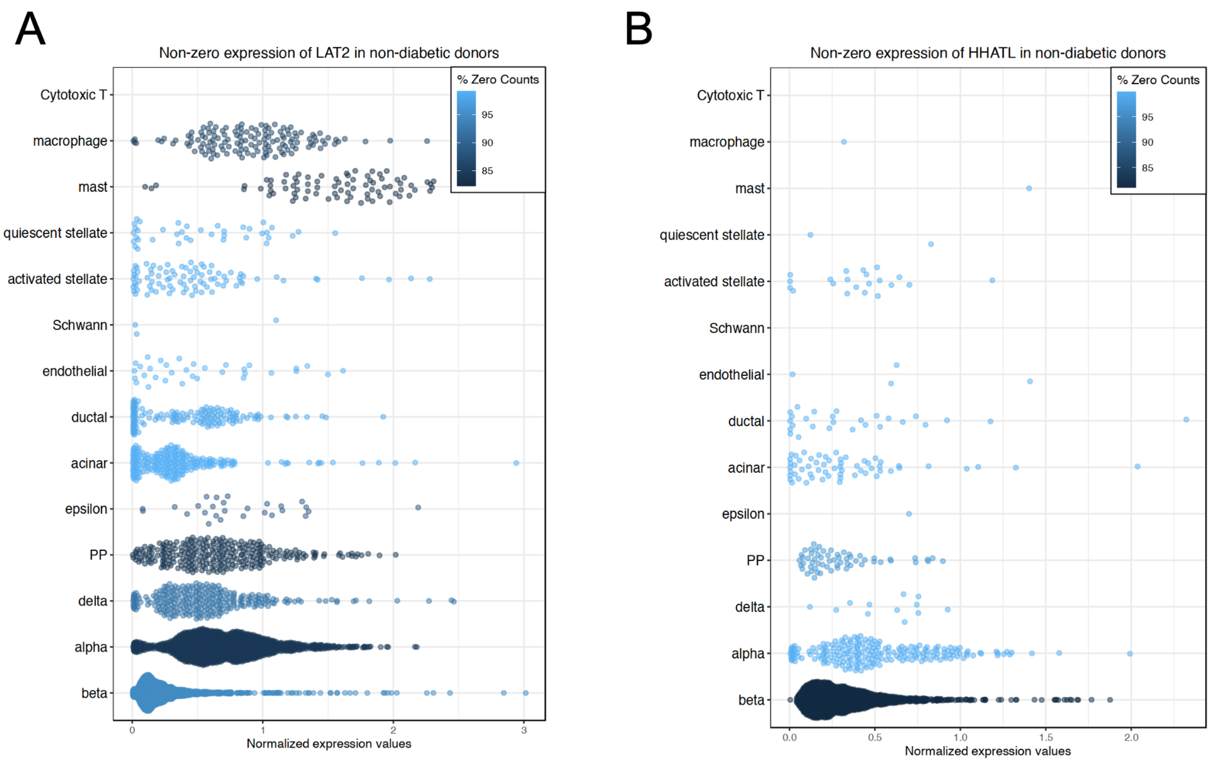


**Suppl Figure 5.** Bulk RNA-seq features with the most significant positive associations with alpha cells (**A**; LAT2, coef = 9.87, FDR = 5.2e-5) and beta cells (**B**; HHATL, coef = 21.20, FDR = 5.8e-7) that were not included as marker genes in the deconvolution analysis.

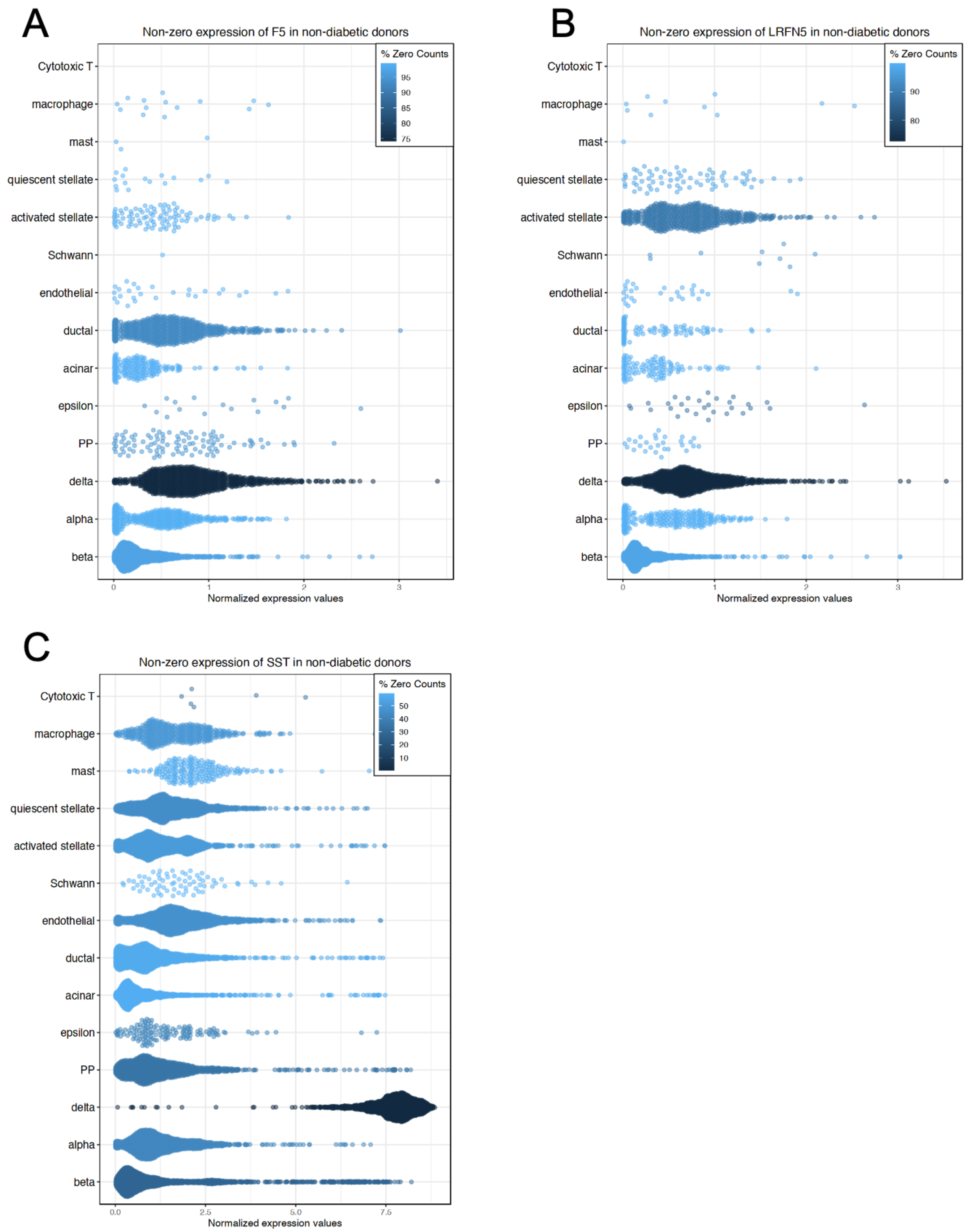


**Suppl Figure 6**. The three transcripts most significantly associated with delta cell proportions in the bulk RNA-seq data: F5 (**A**), LRFN5 (**B**), and SST (**C**).

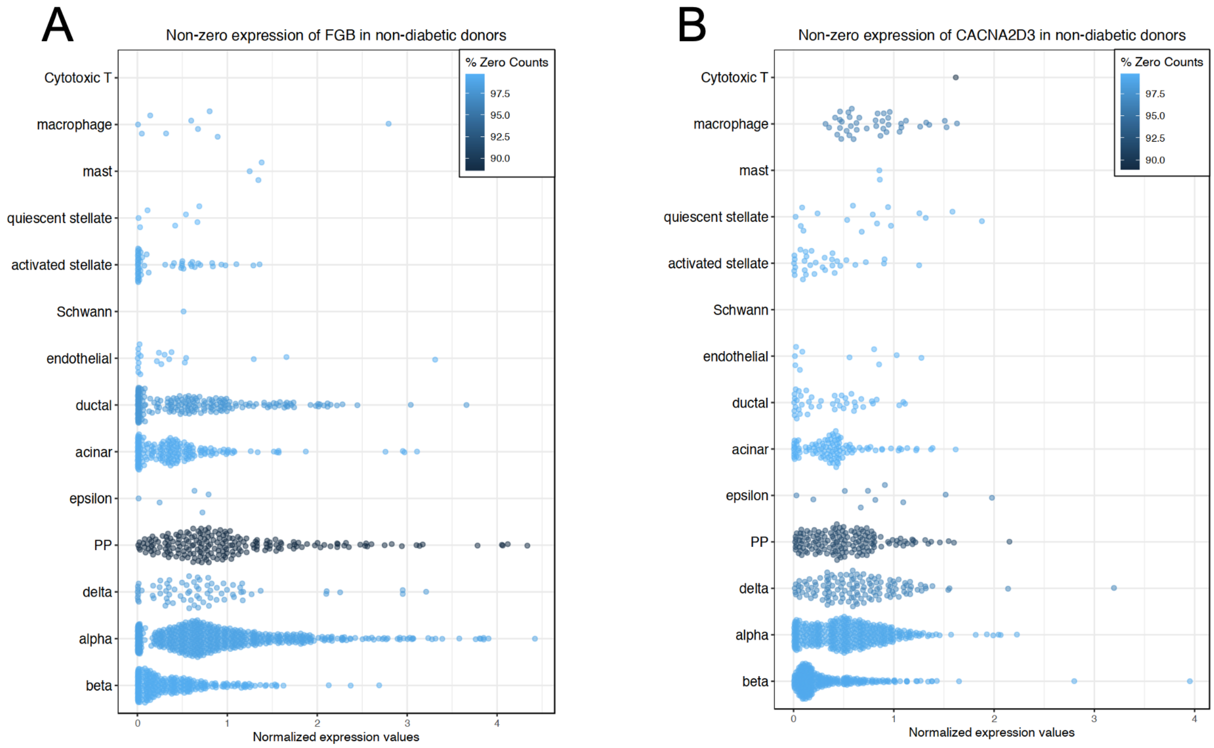


**Suppl Figure 7**. Two of the four transcripts most significantly associated with gamma cell proportions in the bulk RNA-seq data: FGB (**A**; top feature) and CACNA2D3 (**B**; 4th most associated feature).
