## Supplementary Document 1 for "HumanIslets: An integrated platform for human islet data access and analysis"

### HumanIslets Web-Tool

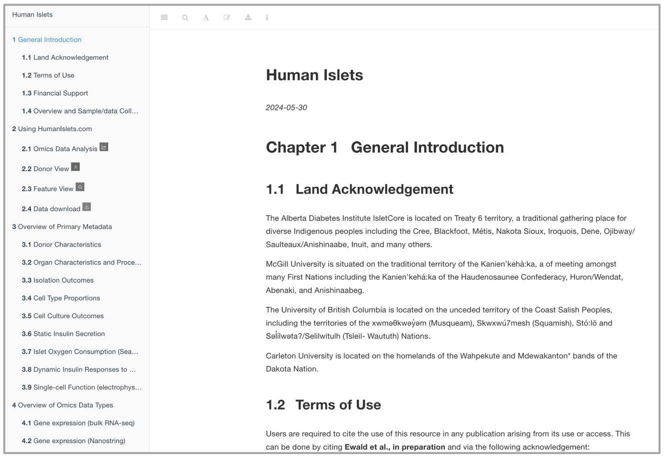


Access extensive documentation, recent tool updates, information on the HumanIslets team and tool roadmap, and Terms of Service here:

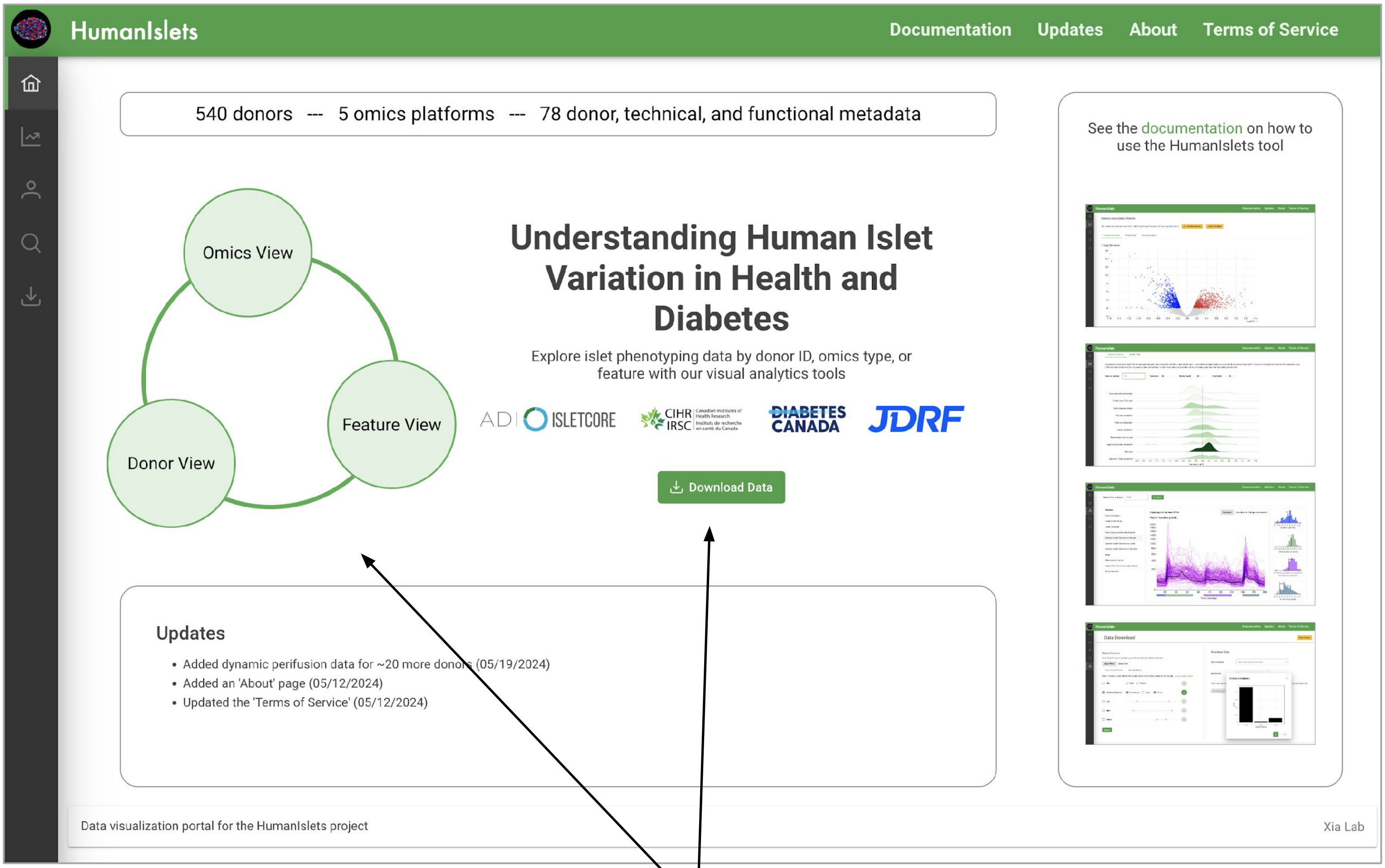


The HumanIslets database can be explored from three complementary views, each accessible from the home page:

1. Omics view: compute gene-level and pathway-level associations between an omics dataset and a metadata variable of interest.
2. Feature view: query all metadata-omics associations that involve a feature of interest, sorted by statistical significance.
3. Donor view: view all metadata and functional outcomes for a specific donor, compared to the distribution of values from other donors.

There is also a data download tool.

### Omics View

The main steps to computing results on this page:

1. Select an omics type of interest (ie. proteomics)
2. Select a metadata or functional outcome of interest. These are grouped into 9 categories. Categorical variables automatically prompt the selection of two specific classes to compare.
3. Select any covariates that you would like to adjust for. The tool supports the inclusion of donor metadata (ie. Age, Sex) or technical variables (ie. cold ischemic time, cell culture, etc).
4. Optionally choose to do the analysis for a subset of donors.
5. Choose a p-value cut-off to determine statistical significance.
6. Click 'Submit'!

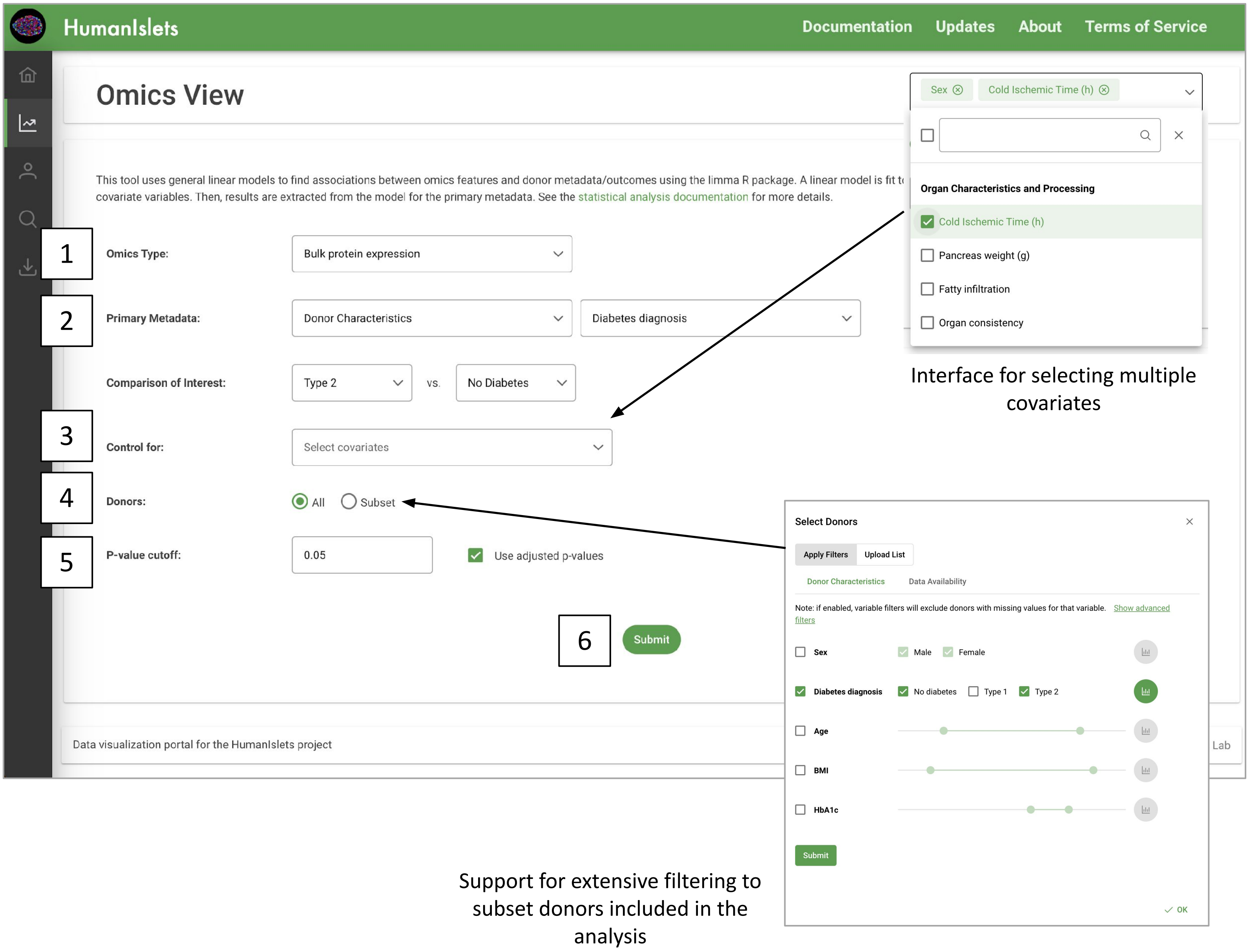

Download date and time-stamped results tables and copies of the input data and R scripts used for the analysis for locally reproducible results.

The results can be explored in three different ways. Here is an introduction to the 'Graphical Summary'.

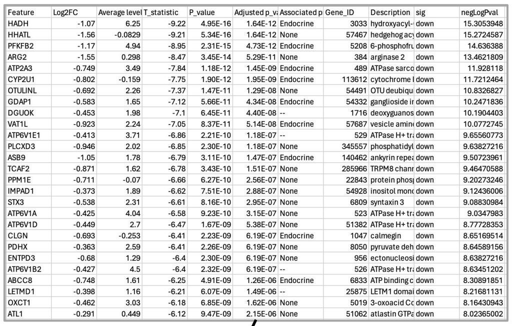

| Feature | Log2FC | Average level | T_statistic | P_value | Adjusted p_va | Associated p | Gene_ID | Description | sig | negLogPval |
|---|---|---|---|---|---|---|---|---|---|---|
| HADH | -1.07 | 6.25 | -9.22 | 4.95E-16 | 1.64E-12 | Endocrine | 3033 | hydroxyacyl- | down | 15.3053948 |
| HHATL | -1.56 | -0.0829 | -9.21 | 5.34E-16 | 1.64E-12 | None | 57467 | hedgehog acy | down | 15.2724587 |
| PFKFB2 | -1.17 | 4.94 | -8.95 | 2.31E-15 | 4.73E-12 | Endocrine | 5208 | 6-phosphofru | down | 14.636388 |
| ARG2 | -1.55 | 0.298 | -8.47 | 3.45E-14 | 5.29E-11 | None | 384 | arginase 2 | down | 13.4621809 |
| ATP2A3 | -0.749 | 3.49 | -7.84 | 1.18E-12 | 1.45E-09 | Endocrine | 489 | ATPase sarco | down | 11.928118 |
| CYP2U1 | -0.802 | -0.159 | -7.75 | 1.90E-12 | 1.95E-09 | Endocrine | 113612 | cytochrome I | down | 11.7212464 |
| OTULINL | -0.692 | 2.26 | -7.37 | 1.47E-11 | 1.29E-08 | None | 54491 | OTU deubiqu | down | 10.8326827 |
| GDAP1 | -0.583 | 1.65 | -7.12 | 5.66E-11 | 4.34E-08 | Endocrine | 54332 | ganglioside ir | down | 10.2471836 |
| DGUOK | -0.453 | 1.98 | -7.1 | 6.45E-11 | 4.40E-08 | -- | 1716 | deoxyguanos | down | 10.1904403 |
| VAT1L | -0.923 | 2.24 | -7.05 | 8.37E-11 | 5.14E-08 | Endocrine | 57687 | vesicle amin | down | 10.0772745 |
| ATP6V1E1 | -0.413 | 3.71 | -6.86 | 2.21E-10 | 1.18E-07 | -- | 529 | ATPase H+ tra | down | 9.65560773 |
| PLCXD3 | -0.946 | 2.02 | -6.85 | 2.30E-10 | 1.18E-07 | None | 345557 | phosphatidyl | down | 9.63827216 |
| ASB9 | -1.05 | 1.78 | -6.79 | 3.11E-10 | 1.47E-07 | Endocrine | 140462 | ankyrin repea | down | 9.50723961 |
| TCAF2 | -0.871 | 1.62 | -6.78 | 3.43E-10 | 1.51E-07 | None | 285966 | TRPM8 chanr | down | 9.46470588 |
| PPM1E | -0.711 | -0.07 | -6.66 | 6.27E-10 | 2.56E-07 | None | 22843 | protein phos | down | 9.20273246 |
| IMPAD1 | -0.373 | 1.89 | -6.62 | 7.51E-10 | 2.88E-07 | None | 54928 | inositol mono | down | 9.12436006 |
| STX3 | -0.538 | 2.31 | -6.61 | 8.16E-10 | 2.95E-07 | None | 6809 | syntaxin 3 | down | 9.08830984 |
| ATP6V1A | -0.425 | 4.04 | -6.58 | 9.23E-10 | 3.15E-07 | None | 523 | ATPase H+ tra | down | 9.0347983 |
| ATP6V1D | -0.449 | 2.7 | -6.47 | 1.67E-09 | 5.38E-07 | None | 51382 | ATPase H+ tra | down | 8.77728353 |
| CLGN | -0.693 | -0.253 | -6.41 | 2.23E-09 | 6.19E-07 | Endocrine | 1047 | calmegin | down | 8.65169514 |
| PDHX | -0.363 | 2.59 | -6.41 | 2.26E-09 | 6.19E-07 | None | 8050 | pyruvate deh | down | 8.64589156 |
| ENTPD3 | -0.68 | 1.29 | -6.4 | 2.30E-09 | 6.19E-07 | None | 956 | ectonucleosi | down | 8.63827216 |
| ATP6V1B2 | -0.427 | 4.5 | -6.4 | 2.32E-09 | 6.19E-07 | -- | 526 | ATPase H+ tra | down | 8.63451202 |
| ABCC8 | -0.748 | 1.61 | -6.25 | 4.91E-09 | 1.26E-06 | Endocrine | 6833 | ATP binding c | down | 8.30891851 |
| LETMD1 | -0.398 | 1.16 | -6.21 | 6.07E-09 | 1.49E-06 | -- | 25875 | LETM1 domai | down | 8.21681131 |
| OXCT1 | -0.462 | 3.03 | -6.18 | 6.85E-09 | 1.62E-06 | None | 5019 | 3-oxoacid Co | down | 8.16430943 |
| ATL1 | -0.291 | 0.449 | -6.12 | 9.47E-09 | 2.15E-06 | None | 51062 | atlastin GTPa | down | 8.02365002 |

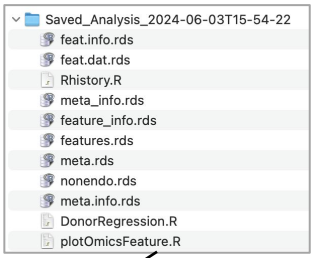


Click a point in the volcano plot to view donor-level measurements in either a violin (categorical metadata) or scatter (continuous metadata) plot

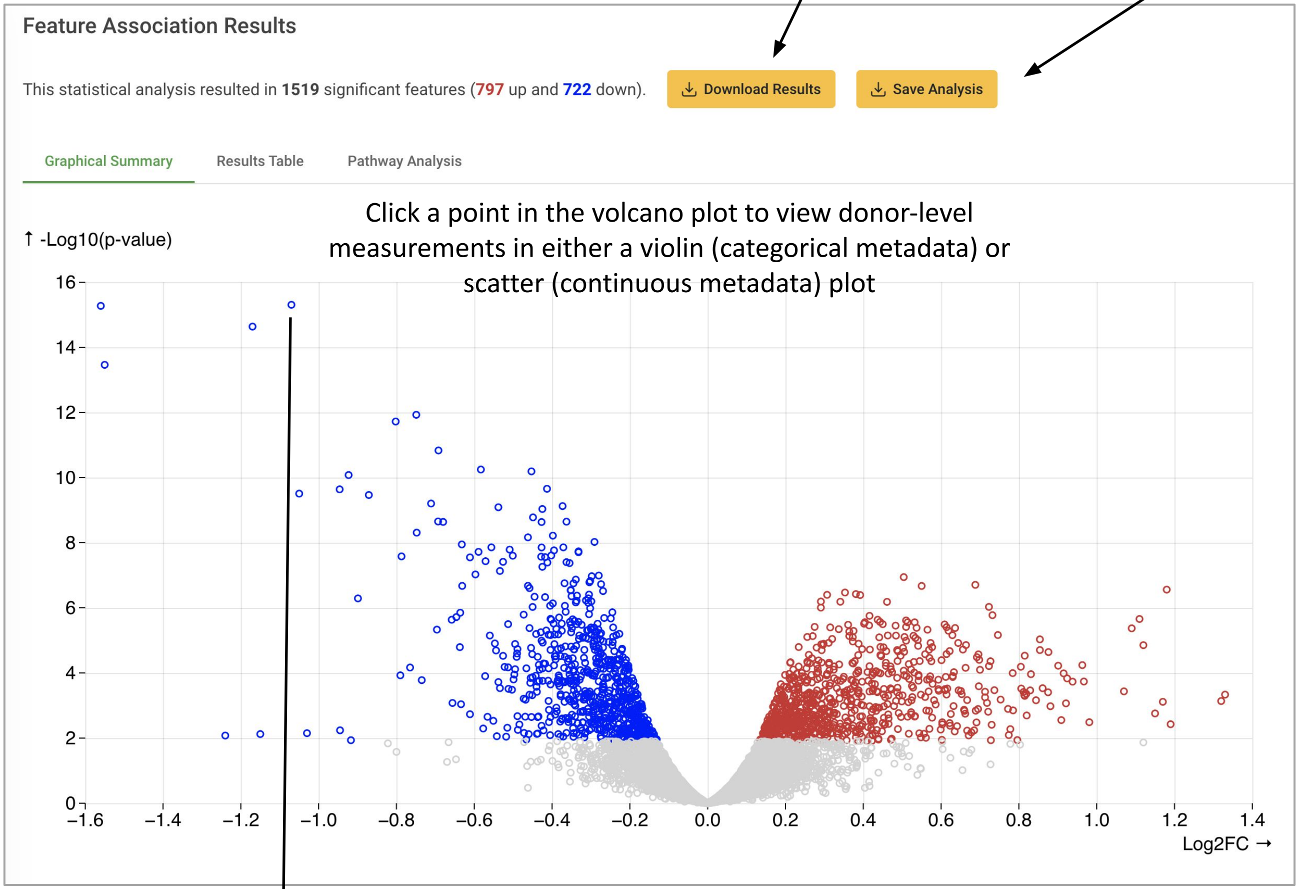


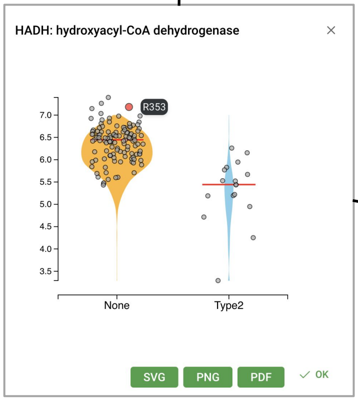


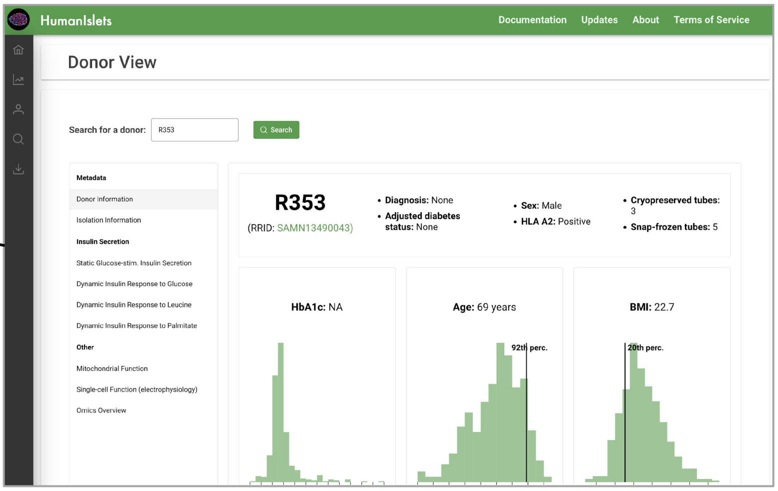


Feature-level plots are interactive. Hovering over a point reveals the donor ID and clicking navigates to the donor-view page for that donor.

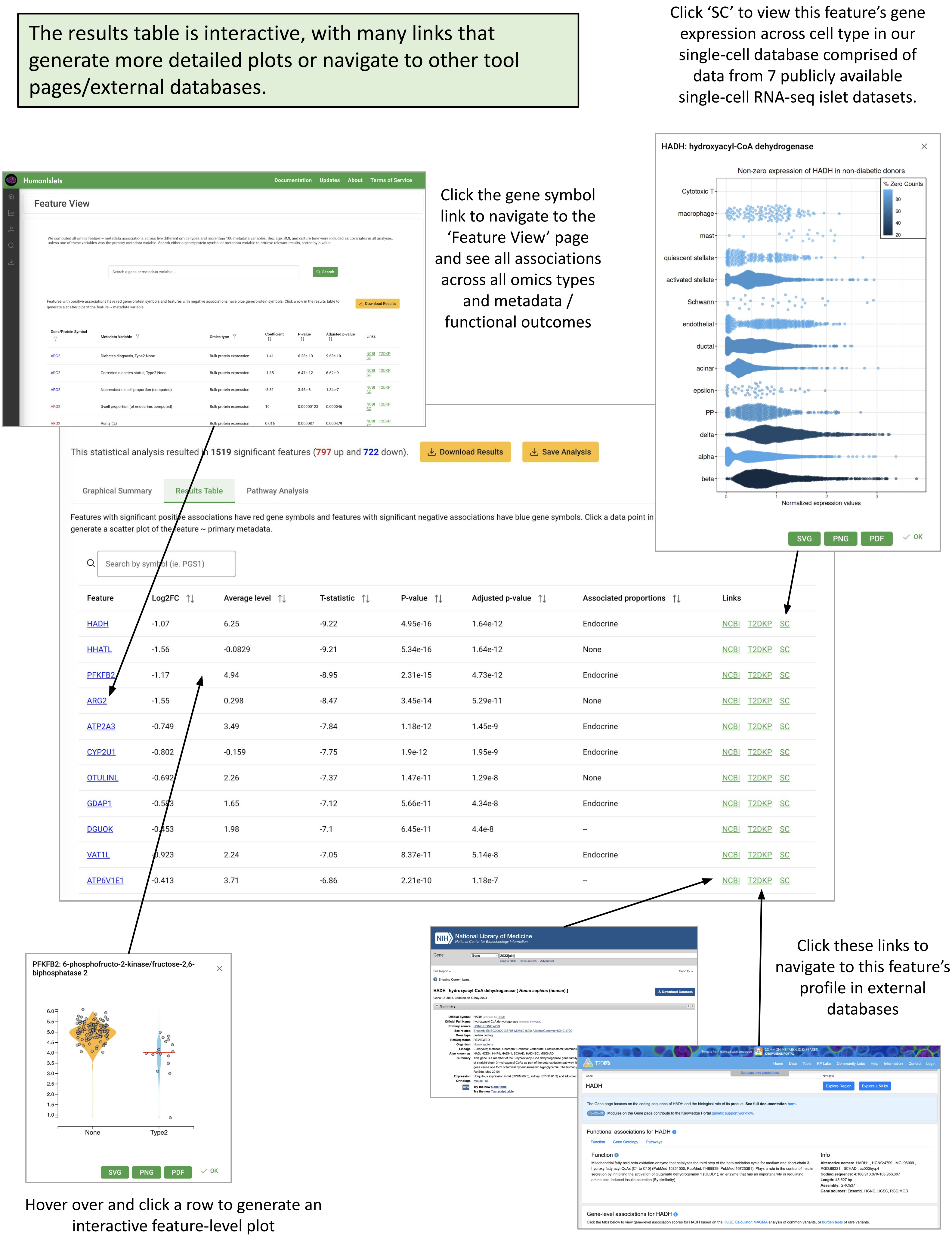

The results table is interactive, with many links that generate more detailed plots or navigate to other tool pages/external databases.
Click 'SC' to view this feature's gene expression across cell type in our single-cell database comprised of data from 7 publicly available single-cell RNA-seq islet datasets.
Click the gene symbol link to navigate to the 'Feature View' page and see all associations across all omics types and metadata / functional outcomes
Hover over and click a row to generate an interactive feature-level plot
Click these links to navigate to this feature's profile in external databases
HumanIslets
Documentation
Updates
About
Terms of Service
Feature View
Download Results
HADH: hydroxyacyl-CoA dehydrogenase
Non-zero expression of HADH in non-diabetic donors
% Zero Counts
Normalized expression values
Cytotoxic T
macrophage
mast
quiescent stellate
activated stellate
Schwann
endothelial
ductal
acinar
epsilon
PP
delta
alpha
beta
SVG
PNG
PDF
OK
This statistical analysis resulted in 1519 significant features (797 up and 722 down).
Download Results
Save Analysis
Graphical Summary
Results Table
Pathway Analysis
Features with significant positive associations have red gene symbols and features with significant negative associations have blue gene symbols. Click a data point in
generate a scatter plot of the feature ~ primary metadata.
Search by symbol (ie. PGS1)
Feature
Log2FC
Average level
T-statistic
P-value
Adjusted p-value
Associated proportions
Links
HADH -1.07 6.25 -9.22 4.95e-16 1.64e-12 Endocrine NCBI T2DKP SC
HHATL -1.56 -0.0829 -9.21 5.34e-16 1.64e-12 None NCBI T2DKP SC
PFKFB2 -1.17 4.94 -8.95 2.31e-15 4.73e-12 Endocrine NCBI T2DKP SC
ARG2 -1.55 0.298 -8.47 3.45e-14 5.29e-11 None NCBI T2DKP SC
ATP2A3 -0.749 3.49 -7.84 1.18e-12 1.45e-9 Endocrine NCBI T2DKP SC
CYP2U1 -0.802 -0.159 -7.75 1.9e-12 1.95e-9 Endocrine NCBI T2DKP SC
OTULINL -0.692 2.26 -7.37 1.47e-11 1.29e-8 None NCBI T2DKP SC
GDAP1 -0.583 1.65 -7.12 5.66e-11 4.34e-8 Endocrine NCBI T2DKP SC
DGUOK -0.453 1.98 -7.1 6.45e-11 4.4e-8 -- NCBI T2DKP SC
VAT1L -0.923 2.24 -7.05 8.37e-11 5.14e-8 Endocrine NCBI T2DKP SC
ATP6V1E1 -0.413 3.71 -6.86 2.21e-10 1.18e-7 -- NCBI T2DKP SC
PFKFB2: 6-phosphofructo-2-kinase/fructose-2,6-biphosphatase 2
None
Type2
National Library of Medicine
HADH hydroxyacyl-CoA dehydrogenase [ Homo sapiens (human) ]
T2DKP
HADH
Functional associations for HADH
Function
Info
Gene-level associations for HADH

The pathway analysis tab supports pathway/gene set analysis of the associations results using two different method (ORA and GSEA) for 8 different gene set libraries.

The ridgeline plot shows the distribution of feature-level fold-changes or t-statistics, colored by the p-value for pathway enrichment. The dimensions of the ridgeline plot can be easily customized to better display results for a specific analysis

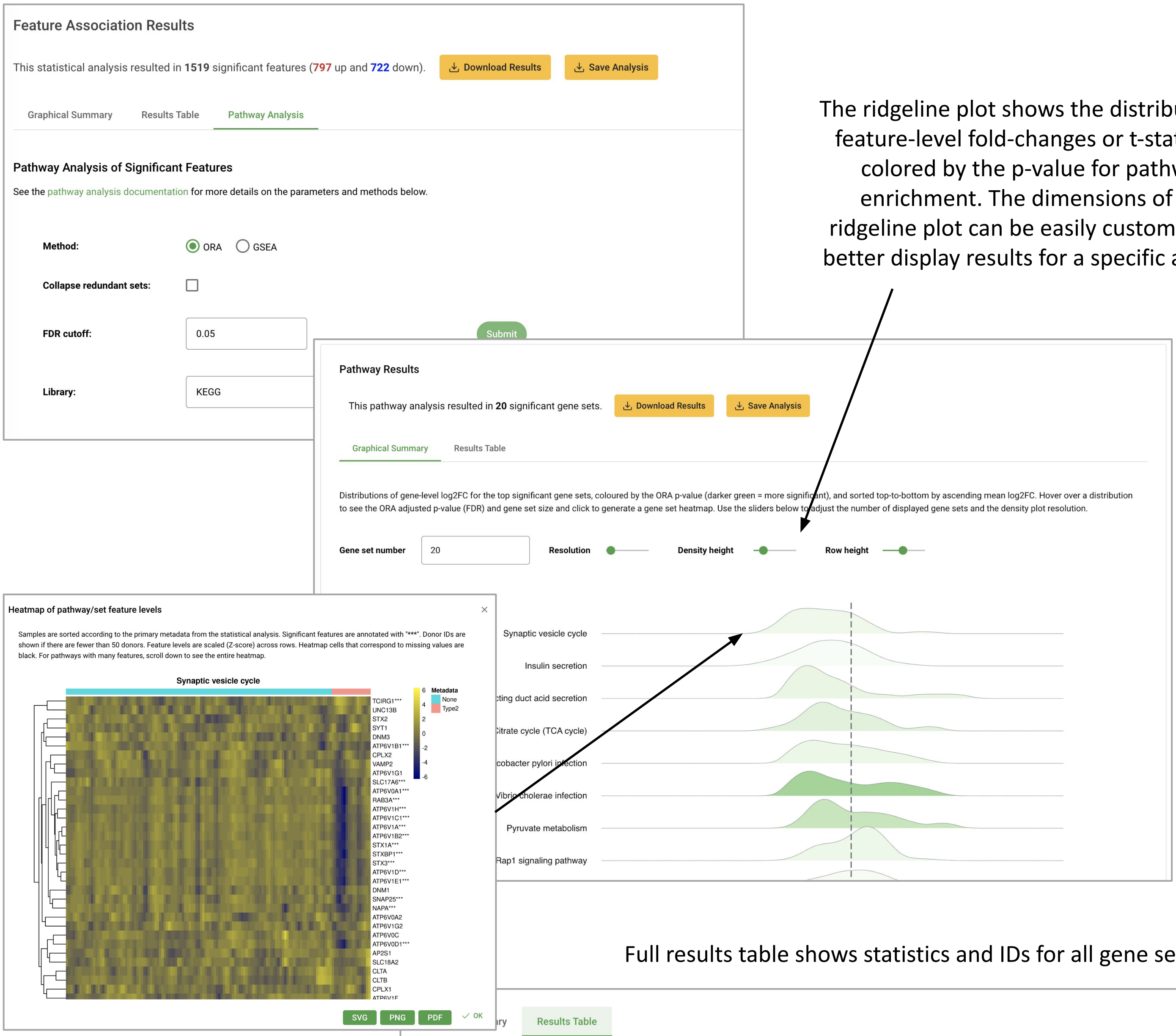


Hover over a pathway result to see the associated statistics, and click to generate a heatmap showing feature-level omics data for that gene set

Full results table shows statistics and IDs for all gene sets

Results Table

| Set Name | P-value | Adj P-value | Hits | Set Size | Set ID |
|---|---|---|---|---|---|
| Ribosome | 3.57e-11 | 8.24e-9 | 62 | 117 | hsa03010 |
| Carbon metabolism | 0.00000195 | 0.000168 | 36 | 71 | hsa01200 |
| Vibrio cholerae infection | 0.00000246 | 0.000168 | 21 | 33 | hsa05110 |
| Biosynthesis of amino acids | 0.00000291 | 0.000168 | 27 | 48 | hsa01230 |
| Glycolysis / Gluconeogenesis | 0.00000535 | 0.000247 | 23 | 39 | hsa00010 |
| Pyruvate metabolism | 0.0000282 | 0.00109 | 17 | 27 | hsa00620 |
| Arginine and proline metabolism | 0.0000554 | 0.00183 | 17 | 28 | hsa00330 |
| Fatty acid degradation | 0.000249 | 0.00656 | 16 | 28 | hsa00071 |
| Citrate cycle (TCA cycle) | 0.000255 | 0.00656 | 11 | 16 | hsa00020 |
| Collecting duct acid secretion | 0.000562 | 0.013 | 11 | 17 | hsa04966 |
| Epithelial cell signaling in Helicobacter pylori infection | 0.00082 | 0.0166 | 17 | 33 | hsa05120 |
| Leukocyte transendothelial migration | 0.000863 | 0.0166 | 24 | 53 | hsa04670 |
| Rap1 signaling pathway | 0.00102 | 0.0181 | 34 | 84 | hsa04015 |

### Feature View

Statistical associations between each omics type and each metadata/function outcome were computed and stored in a database (>3.8 million relationships). Age, sex, BMI, and culture time were included as covariates in the analysis. The pre-computed results can be queried by either gene symbol or by metadata/functional outcome name.

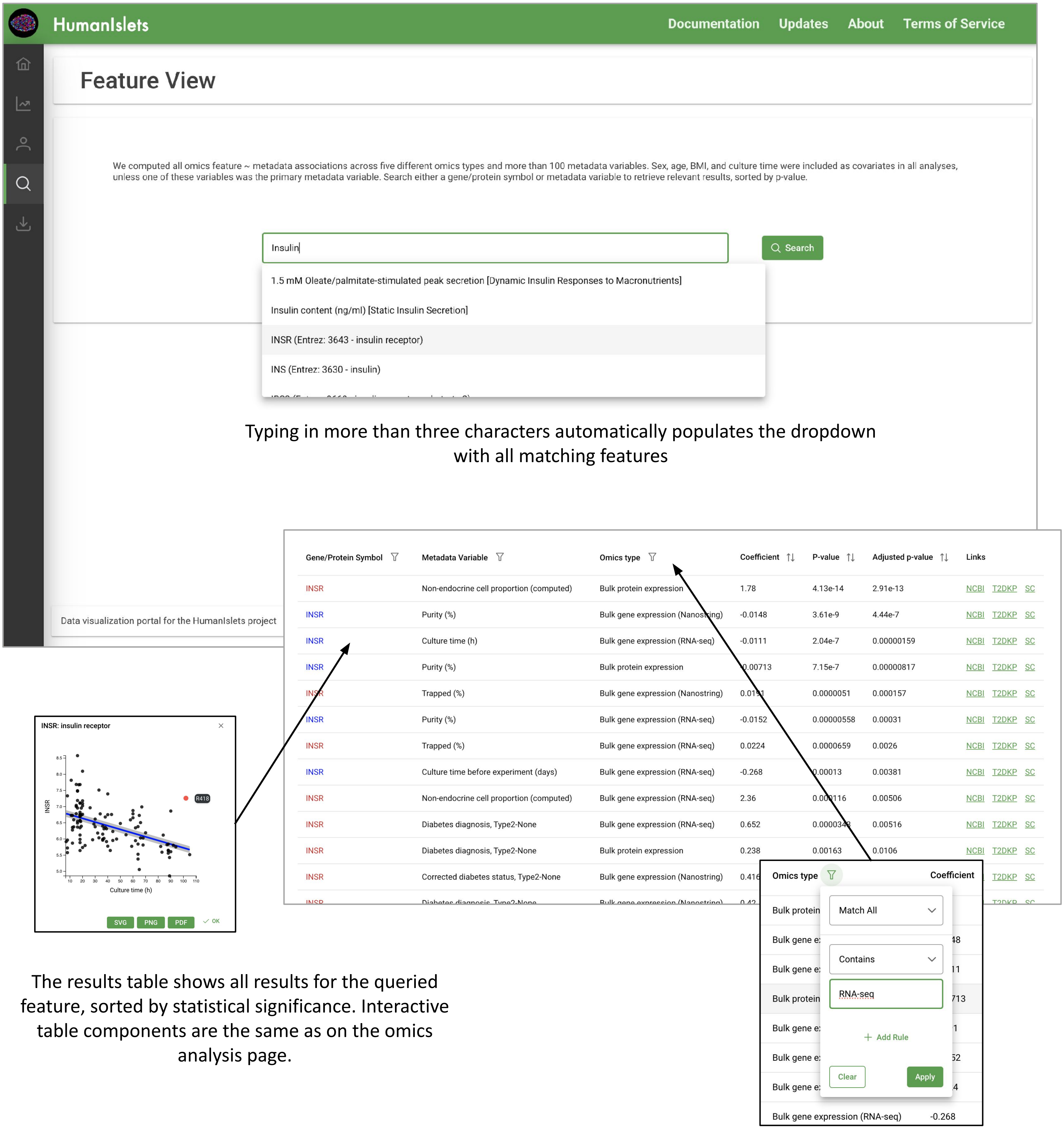


Typing in more than three characters automatically populates the dropdown with all matching features

The results table shows all results for the queried feature, sorted by statistical significance. Interactive table components are the same as on the omics analysis page.

The results table enables filtering, for example by omics type

### Donor View

The Donor View page shows a summary of each donor's outcomes, displayed against the distribution of values from different donors across the HumanIslets database. Donors can be searched by either donor ID (ie. R360) or RRID (ie. SAMN14146088).

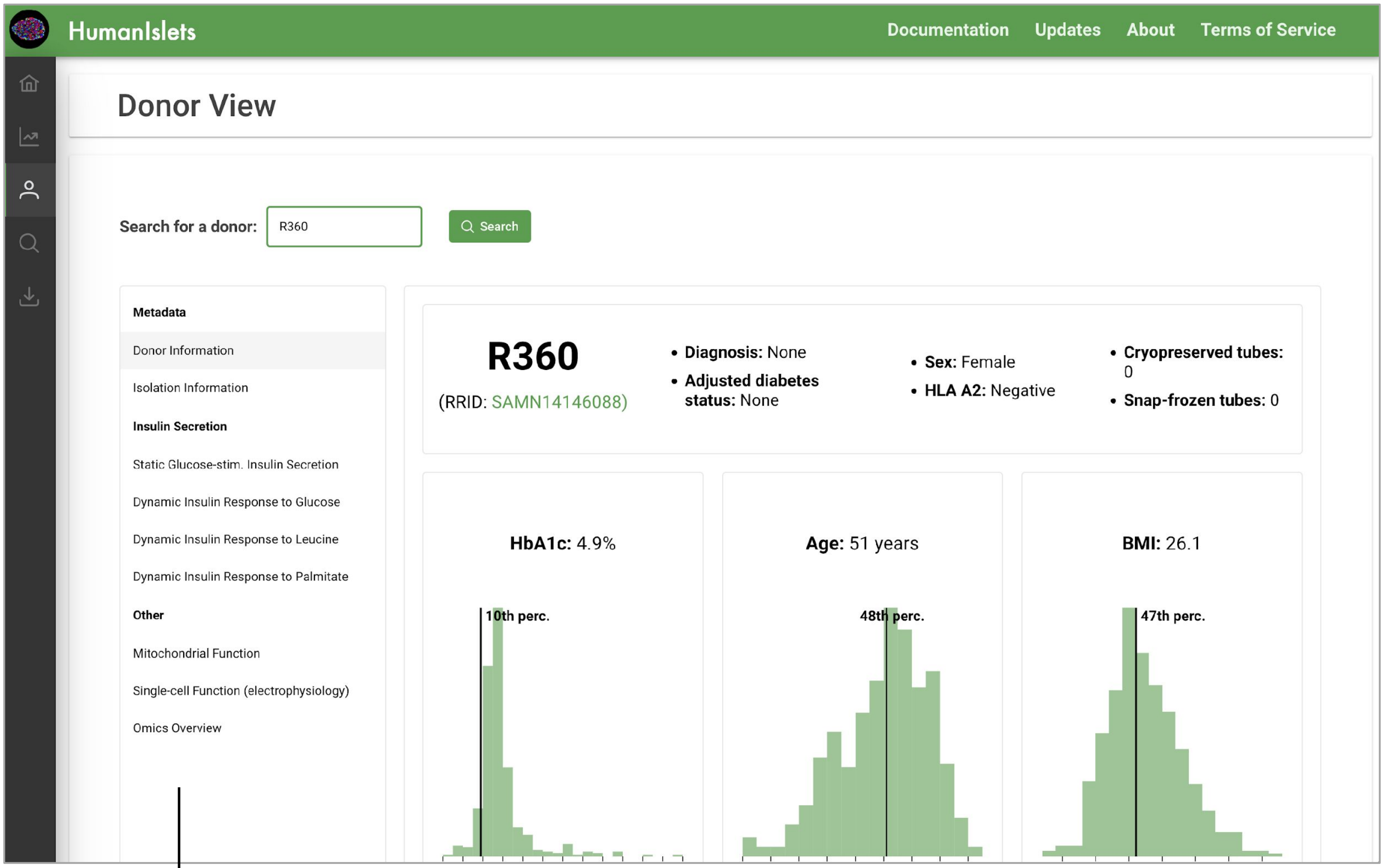


There are currently nine types of data available to view. More data will be added over time as it is collected.

Each tab shows a summary of a different kind of dataset. For example …

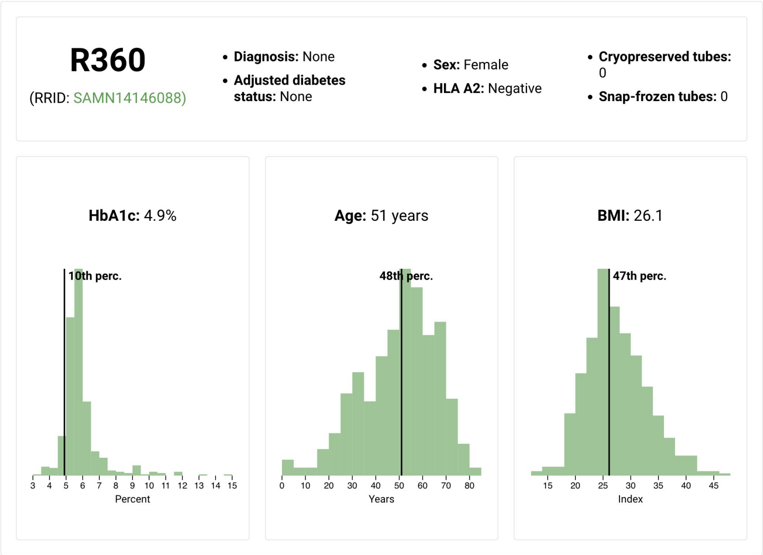


Basic donor metadata, including availability of isolated islets for future research projects

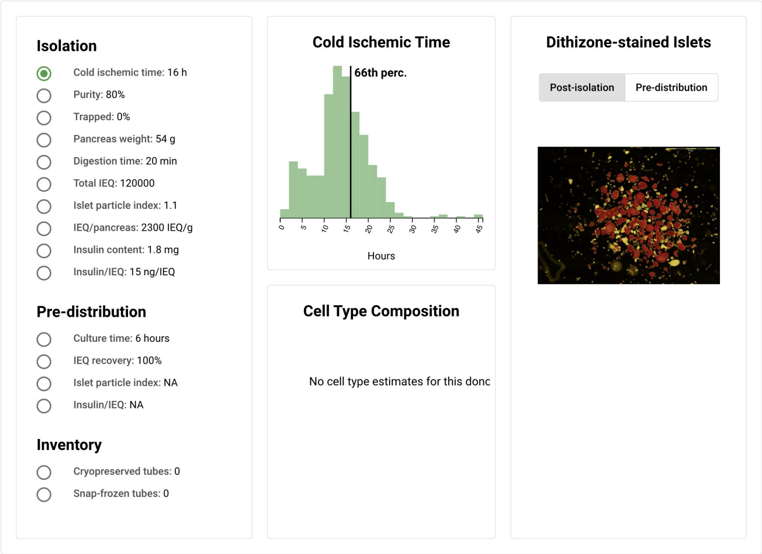


Technical islet isolation outcomes, including images of the isolated islets

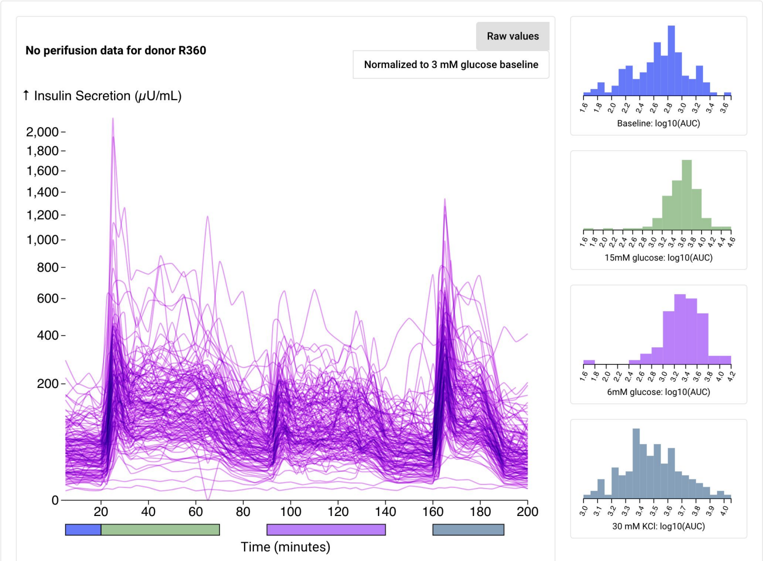


Functional outcomes such as dynamic insulin secretion in response to three different nutrients. Functional data also include static insulin secretion, oxygen consumption and single-cell electrophysiology outcomes.

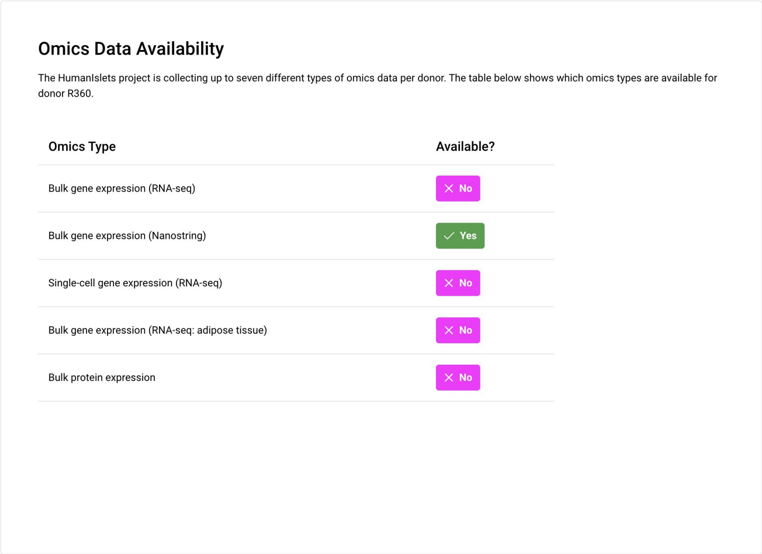


An overview of the omics data available for this donor

### Data Download

The data download page has two steps: ‘Select Donors’ and ‘Download Data’. In the first step, users can filter donors by metadata, data availability, or a donor list.

Clicking this generates a distribution of the values, helping users choose a cut-off

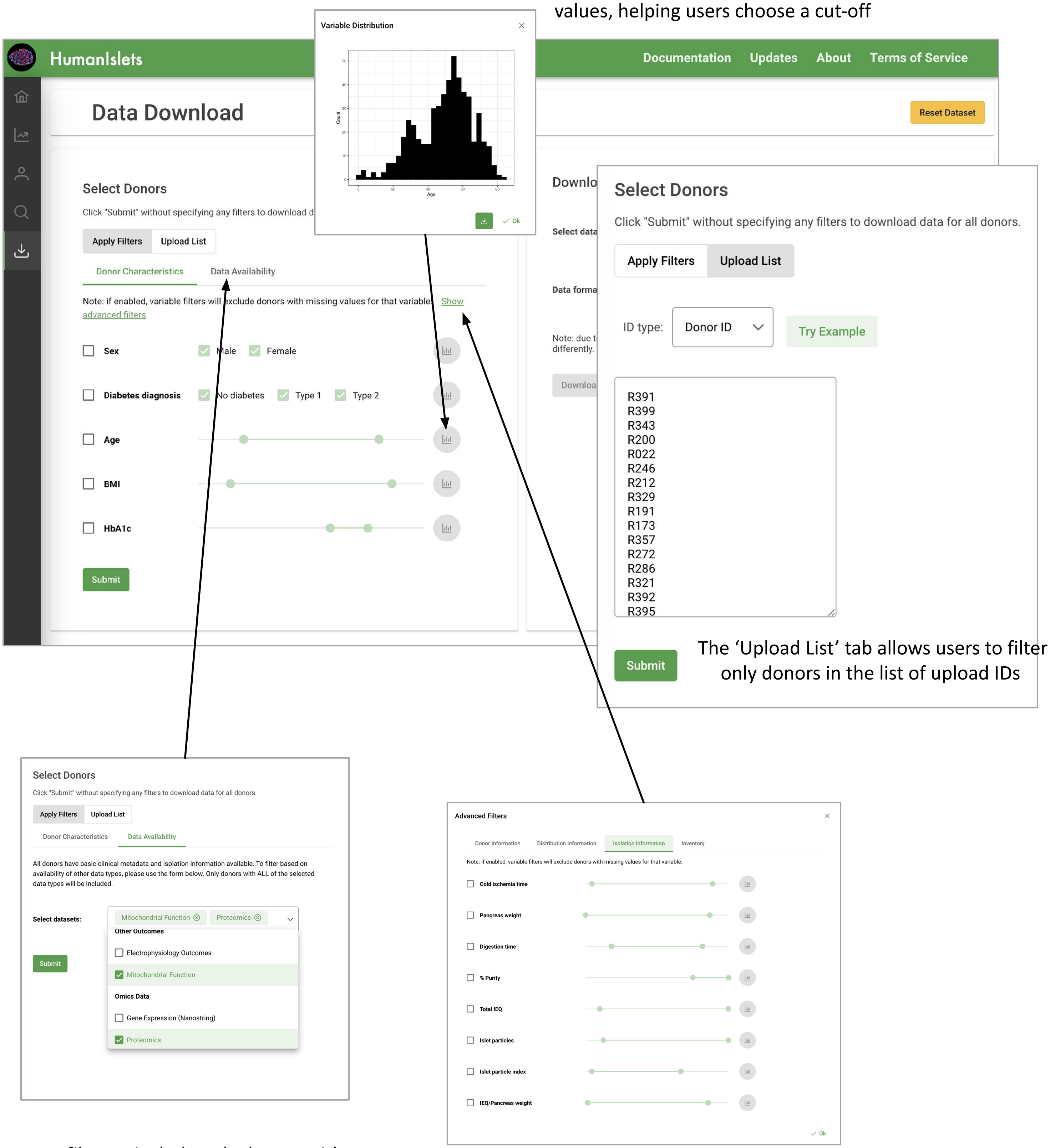


Users can filter to include only donors with specific data types available

Clicking the advanced filters link allows users to filter by many more variables

Once the donors have been selected, users can choose the data types that they'd like to download and the format that they want it in.

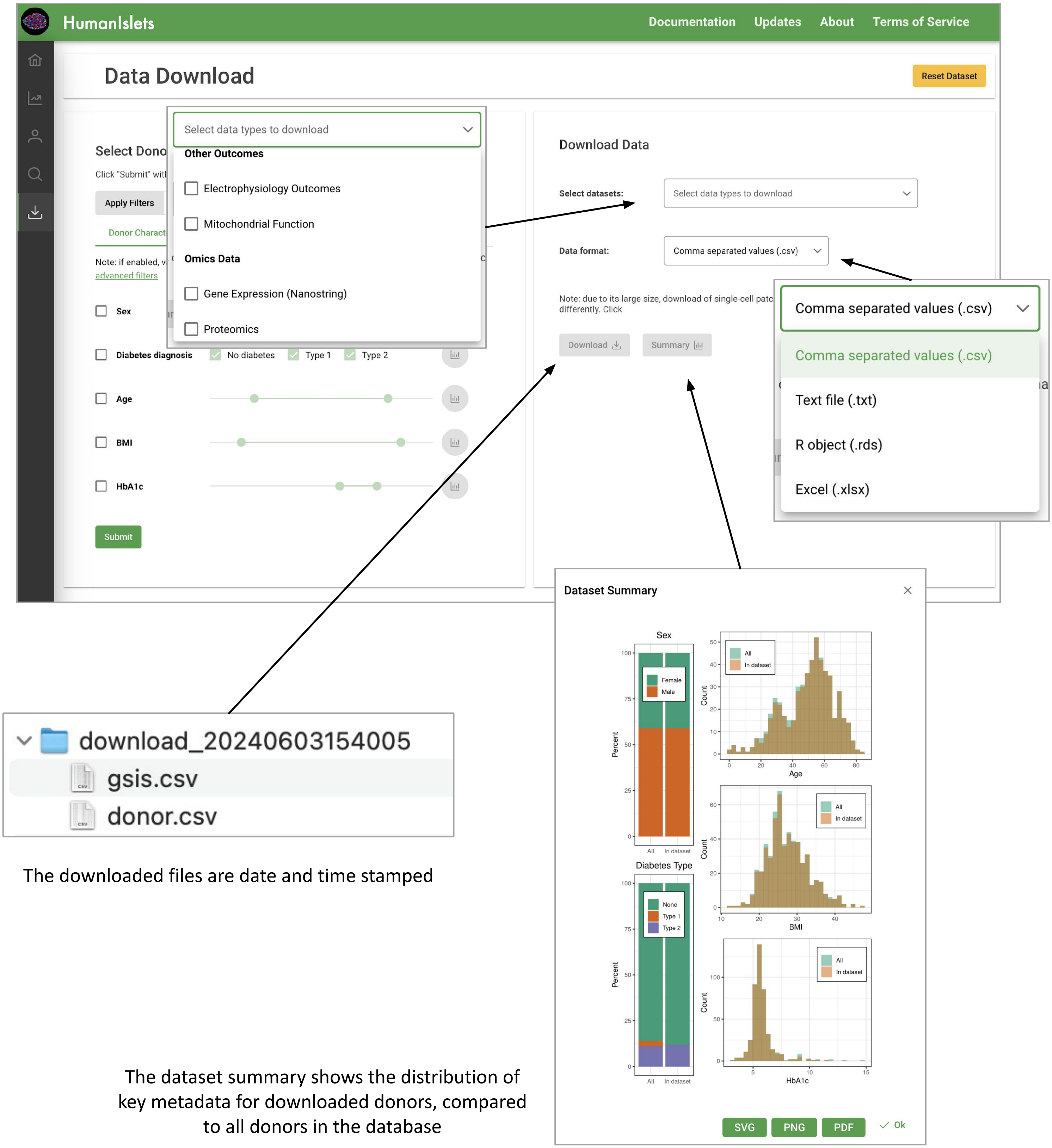


The downloaded files are date and time stamped

The dataset summary shows the distribution of key metadata for downloaded donors, compared to all donors in the database
