## Supplementary Document 2 for "HumanIslets: An integrated platform for human islet data access and analysis"

### Ch 1 - General Introduction

##### 1.1 Land Acknowledgement

The Alberta Diabetes Institute IsletCore is located on Treaty 6 territory, a traditional gathering place for diverse Indigenous peoples including the Cree, Blackfoot, Métis, Nakota Sioux, Iroquois, Dene, Ojibway/ Saulteaux/Anishinaabe, Inuit, and many others.

McGill University is situated on the traditional territory of the Kanien’kehà:ka, a of meeting amongst many First Nations including the Kanien’kehá:ka of the Haudenosaunee Confederacy, Huron/Wendat, Abenaki, and Anishinaabeg.

The University of British Columbia is located on the unceded territory of the Coast Salish Peoples, including the territories of the xwməθkwəy̓əm (Musqueam), Skwxwú7mesh (Squamish), Stó:lō and Səl̓ílwətaʔ/Selilwitulh (Tsleil- Waututh) Nations.

Carleton University is located on the homelands of the Wahpekute and Mdewakanton* bands of the Dakota Nation.

##### 1.2 Terms of Service

Users are required to cite the use of this resource in any publication arising from its use or access. This can be done by ***citing ‘Ewald et al., in preparation’*** and via the following acknowledgement:

> ***“This work includes data and/or analyses from HumanIslets.com funded by the Canadian Institutes of Health Research, JDRF Canada, and Diabetes Canada (5-SRA-2021-1149-S-B/TG 179092) with data from islets isolated by the Alberta Diabetes Institute IsletCore with the support of the Human Organ Procurement and Exchange program, Trillium Gift of Life Network, BC Transplant, Quebec Transplant, and other Canadian organ procurement organizations with written informed donor consent as approved by the Human Research Ethics Board at the University of Alberta (Pro00013094).”***

The HumanIslets tool relies on statistical methods and R packages published by others. We encourage researchers to cite these publications in addition to our manuscript, based on the citations and links provided throughout the tool interface, manuscript methods, and in this Documentation resource.

Data and analysis tools are provided without warranty, are not for clinical use, and should not be used directly or indirectly to inform clinical diagnoses or treatment. While every effort has been made to ensure the accuracy, integrity, and validity of the data and analyses presented here for research purposes only, users are solely responsible for any results and interpretations.

Because datasets are occasionally updated, which may impact normalisation and analyses, ***users should report the date of web tool access, analysis, and/or data download***. To facilitate this, data downloads are provided with a time/date stamp.

#### 1.3 Financial Support

Data collection has been supported by research grants from the Canadian Institutes of Health Research (MacDonald - 186226, 148451; Johnson - 168857), JDRF (MacDonald - 2-SRA-2019-698-S-B; Pepper - 5-CDA-2020-945-A-N), BCCHRI Child Health Integrative Partnership Strategy Funding (Verchere, Levings and Lynn), the National Institutes of Health (MacDonald - U01-DK-120447; MacDonald/Gloyn - U01-DK-123716; Gloyn - U01-DK105535, U01-DK085545, UM-1DK126185), and the Wellcome Trust (Gloyn - 095101, 200837, 098381, 106130, 203141, 203141) and the National Institute for Health Research (NIHR) Oxford Biomedical Research Centre (BRC). Proteomics infrastructure and analysis was supported by the UBC Life Sciences 50 Institute, Canada Foundation for Innovation, BC Knowledge Development Fund, and Genome Canada/BC (PRO264). Pepper holds the Canada Research Chair (CRC) in Cell Therapies for Diabetes (CRC-2021-0026) and MacDonald holds the CRC in Islet Biology (CRC-2019-00334).

Some data used in this web tool includes patch-seq data, and single-cell RNAseq used for cell type expression analysis, from the Human Pancreas Analysis Program (HPAP-RRID:SCR_016202) Database (https://hpap.pmacs.upenn.edu), a Human Islet Research Network (RRID:SCR_014393) consortium (UC4-DK-112217, U01-DK-123594, UC4-DK-112232, and U01-DK-123716).

Consolidation of datasets and web tool development was supported by a research grant funded by the Canadian Institutes of Health Research, JDRF Canada, and Diabetes Canada (5-SRA-2021-1149-S-B/TG 179092) to MacDonald (Alberta), Xia (McGill), Johnson (UBC) and Bruin (Carleton) with collaborators Gloyn (Stanford), Foster (UBC), Atlas (Ottawa), Mulvihill (Ottawa), Lynn (UBC), and Verchere (UBC).

#### 1.4 Overview and Sample/data Collection

The purpose of the HumanIslets.com tool is to allow exploration and analysis of, and access to data from, human islet molecular and functional phenotyping studies. We hope this resource will facilitate research and hypothesis generation in a manner that is accessible to both experts and non-experts in data analysis. The **below figure** illustrates general workflows, samples, and datasets collected as part of this initiative Data noted in blue are currently included within the web tool, while those in grey are are planned for addition at a later date.

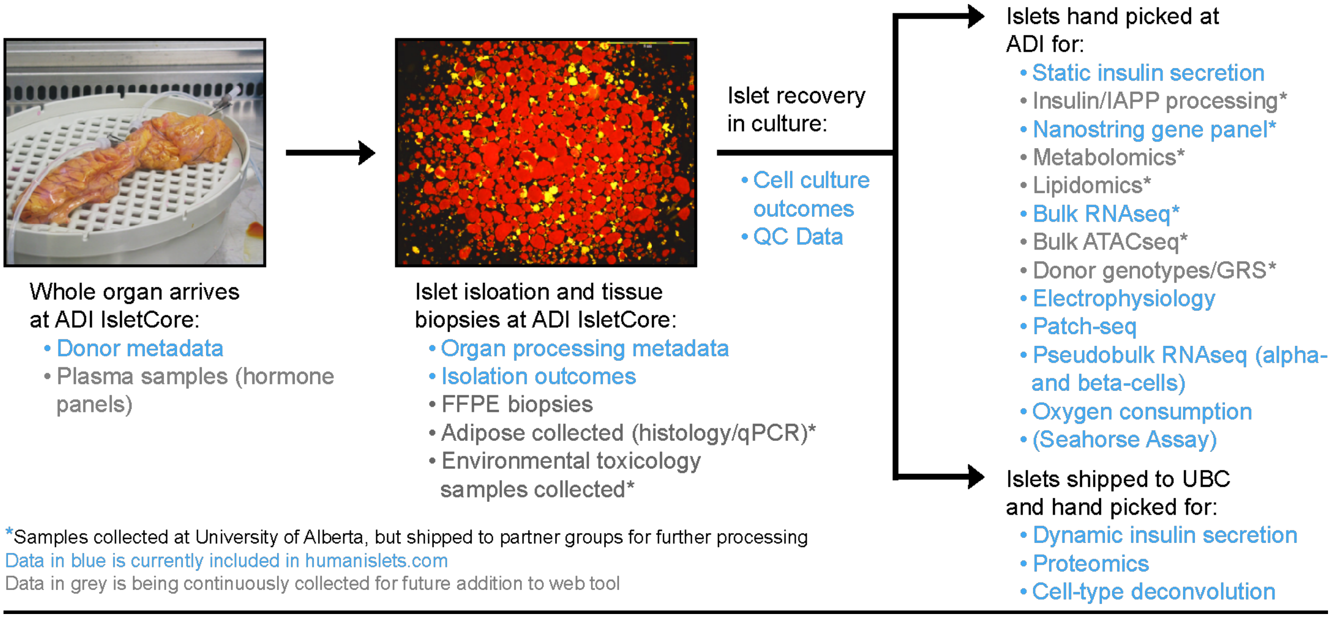
Whole organ arrives
at ADI IsletCore:
• Donor metadata
• Plasma samples (hormone panels)
Islet isloation and tissue
biopsies at ADI IsletCore:
• Organ processing metadata
• Isolation outcomes
• FFPE biopsies
• Adipose collected (histology/qPCR)*
• Environmental toxicology samples collected*
Islet recovery
in culture:
• Cell culture outcomes
• QC Data
Islets hand picked at
ADI for:
• Static insulin secretion
• Insulin/IAPP processing*
• Nanostring gene panel*
• Metabolomics*
• Lipidomics*
• Bulk RNAseq*
• Bulk ATACseq*
• Donor genotypes/GRS*
• Electrophysiology
• Patch-seq
• Pseudobulk RNAseq (alpha- and beta-cells)
• Oxygen consumption
• (Seahorse Assay)
Islets shipped to UBC
and hand picked for:
• Dynamic insulin secretion
• Proteomics
• Cell-type deconvolution
*Samples collected at University of Alberta, but shipped to partner groups for further processing
Data in blue is currently included in humanislets.com
Data in grey is being continuously collected for future addition to web tool

### Ch 2 - Using HumanIslets.com

##### 2.1 Omics View 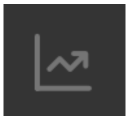

HumanIslet.com presently includes information from 540 human research islet isolations that includes donor and technical metadata, islet and islet cell function, and omics. Omics datasets include bulk proteomics, bulk RNAseq, gene panels (Nanostring), patch-seq, and pseudobulk RNAseq for alpha- and beta-cells. Omics-metadata relationships can be analysed by:

1. Selecting the 'Omics Type' of interest
2. Selecting the 'Primary Metadata' to be analysed, and 'Comparison of Interest' if applicable
3. Choosing covariates to control for
4. Selecting donors to include
    a. Choosing 'Subset' will open a window that allows for filtering by donor characteristics and data availability, or upload of a list of donor IDs or RRIDs of interest
5. Selecting a p-value cutoff, and the use of adjusted p-values
6. Click 'submit'

The number of donors included in the analysis will be displayed. If too few donors are available for the chosen analysis (<10), an error message will be shown. Results of the analysis will be shown under 'Feature Association Results', and both results and analysis pipeline can be downloaded by clicking on the appropriate button. Results are shown as a graphical representation (volcano plot) or table, accessible by clicking on the appropriate tab. Clicking on features (points in the volcano plot, or table entries) will raise a graph showing categorical or scatterplot representations of the data/feature of interest. Clicking on individual donors within those plots will take you to the 'Donor View' page for that donor (section 2.2, below), allowing for inspection of potential outlier donors. In the Results Table, clicking on the feature name will take you to the 'Feature View' page (section 2.3, below) for that gene/protein. At the right are links to the NCBI gene page and T2D Knowledge Portal (T2DKP) entries for each feature (external sites), and to a single-cell RNAseq reference dataset (SC) to inspect cell-type specific expression of the feature.

The third tab under the 'Feature Association Results' pulls up a Pathway Analysis tool for common Overrepresentation Analysis (ORA) or Gene Set Enrichment Analysis (GSEA) on the results. To perform pathway analysis:

1. Select ORA or GSEA
    a. ORA is performed on significant hits based on the p-value threshold set above

    b. GSEA can be performed using log2FC/coefficient or T-statistic for ranking, and can be performed even in cases when no individual features are marked as significant
2. Select whether to collapse redundant pathways
3. Choose a false discovery rate (FDR) cutoff
4. Choose among common pathway libraries to query
5. Click ‘submit’

The ‘Pathway Results’ will be displayed in both graphical and table format, selected by clicking on the appropriate tab. As above, both results and analysis pipeline can be downloaded by clicking on the appropriate button. In the interactive graphical view, significant pathways (based on FDR cutoff) are shown as ridgeplots. Clicking on these will raise heatmaps showing the contribution of individual genes/proteins.

**2.2 Donor View** 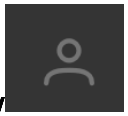

The Donor View page allows examination of individual donor, islet isolation, and functional data from each of 540 donors currently in the HumanIslets.com database. Donors can be searched by ADI IsletCore Donor ID or Research Resource ID (RRID). Data categories can be selected in tabs indicated on the left (Donor information; Isolation Information; Static and Dynamic Insulin Secretion; Mitochondrial Function; Electrophysiology, along with an overview of Omics data currently available). These metadata are described in detail in chapters 3 and 4, below.

Typically, data is shown for selected donor in the context of the entire dataset, allowing for easy inspection and determination of whether a donor falls within the normal range of the population. A few points that are worth noting here:$

- *Donor Information* shows the RRID if available. Clicking on this will link out to the NCBI Biosample page for this donor
- *Isolation Information* also contains the current inventory of snap frozen and cryopreserved islet tubes for that donor (all donors also have paraffin embedded biopsies)
- *Isolation Information* includes cell type composition calculated by deconvolution in donors with bulk proteomics data (see section 5.5, below)
- *Dynamic Insulin Responses* can be shown with or without normalisation to baseline
- *Mitochondrial Function* (oxygen consumption by Seahorse assay) can be shown with or without normalisation to DNA and baseline

**2.3 Feature View** 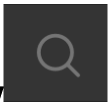

Instead of performing an ‘Omics Analysis’ focused on a particular omics type and metadata variable, users can search for a feature of interest and pull up ALL associations for that particular feature of interest, corrected for common covariates (see section 5.4, below). Searchable features include genes/proteins and also metadata variable. This allows rapid interrogation of precomputed results across all datasets.

Searching for a gene/protein feature generates a table of results ranked by adjusted p-value. Searching for a metadata variable (e.g. BMI) generates a table of feature (gene/protein) results across all omics datasets, ranked by adjusted p-value. Clicking on rows and external links brings up categorical or scatterplots of the association, external pages for the feature of interest, or single-cell RNAseq expression for that feature. Because a large number of associations can be displayed, filtering of the metadata or omics type is enabled by clicking on the column title.

##### 2.4. Data Download 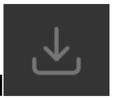

All data at HumanIslets.com is available for download. Donors of interest can be filtered by Donor Characteristics (see section 3.1) and/or data availability. Advanced filters allow selection of donors of interest by any metadata available. Donors of interest can also be selected by uploading a list of donor IDs (for example, donor IDs obtained from a particular Omics Analysis). Data can be downloaded as Comma separated values (.csv), text (.txt), R object (.rds), or in Excel format (.xlsx) within a zip file. To perform data download:

1. Under ‘Select Donors’ choose filtering parameters, upload a list of donor IDs
    a. Distribution of chosen donors compared with the entire dataset can be assessed by clicking on the icons at right
2. Once donors of interest are selected/filtered, click ‘submit’ to choose those donors
    a. The total number of donors chosen will be displayed
3. Under ‘Download Data’
    a. Select the datasets of interest
    b. Select the data format
    c. The distribution of datasets selected, compared with the entire database, can be shown by clicking ‘Summary’
4. Click ‘Download’

#### Ch 3 - Overview of Primary Metadata

##### 3.1 Donor Characteristics

De-identified donor characteristics, including HLA-typing, collected at the time of organ procurement are provided in a de-identified manner to the ADI IsletCore team by the organ procurement organisations. Users should be aware that some information, such as diabetes diagnosis, may not be clinically validated at the time of organ retrieval. Whenever possible,

HbA1c measurements are performed directly by the ADI IsletCore using blood samples sent with the organ, and donors indicated as type 1 diabetes (T1D) are validated as much as possible through genetic testing, autoantibody testing, and immunohistochemical analysis. While the Donor Metadata ***'diabetes diagnosis'*** reflects the status indicated by the organ procurement organization, the Donor Metadata ***'corrected diabetes status'*** indicates a classification based on the totality of information available to the ADI IsletCore (including 'pre-diabetes' defined as HbA1c between 5.7 and 6.5%, and 'non-diagnosed' donors with HbA1c>6.5% as type 2 diabetes (T2D)).

> *Available donor characteristics metadata include:* Age, Sex, Donation Type (NDD, DCD, MAID), BMI, HbA1c (%), HLA A2 (Yes/No), Diabetes diagnosis, and Corrected diabetes status.

##### 3.2 Organ Characteristics and Processing

Organ characteristic and processing metadata is recorded during the islet isolation by the ADI IsletCore lead technician. Cold ischemia time is calculated from the cross clamp time during organ procurement until the start of the isolation in Edmonton. Please note that some measures, such as 'organ consistency' are subjective.

> *Available organ characteristics and processing metadata include:* Cold Ischemic Time (h), Pancreas weight (g), Fatty infiltration (Yes/No), Organ consistency (Soft, Normal, Fibrotic, Inconsistent), Collagenase supplier (Roche, VitaCyte, Serva, Sigma), Colleganse type (Liberase, Clzyme, Gold, NB1, Sigma, Other, Recombinant), Digestion time (min).

##### 3.3 Isolation Outcomes

Islet purity, islet equivalents (IEQ), % trapped, and islet particle index (IPI) are determined by the ADI IsletCore lead technician. Full details can be found here or in our IsletCore welcome booklet. Details on determining Insulin and DNA samples as part of our quality control can be found here.

> *Available isolation outcomes metadata include:* Total islet equivalents (IEQ), Islet equivalents per pancreas weight (IEQ/g), Purity (%), Trapped (%), Islet particle index, Insulin content (ug), DNA content (ug), Insulin:DNA ratio, Insulin per IEQ (ng/IEQ).

##### 3.4 Cell Type Proportions

Cell proportions are estimated based on deconvolution of bulk islet proteomic data from hand-picked islets using canonical proteins as markers (see below). Estimated cell type proportions are shown as either the 'non-endocrine cell proportion' (of all cells) or as the proportion of the endocrine compartment only.

*Available cell type proportions metadata include:* Non-endocrine cell proportion (computed), beta-cell proportion (of endocrine; computed), alpha-cell proportion (of endocrine; computed), delta-cell proportion (of endocrine; computed), and gamma-cell proportion (of endocrine; computed).

##### 3.5 Cell Culture Outcomes

Following isolation, islets are cultured prior to shipment and sample collection for assay. On the day of shipping/collection, islets are recounted to determine islet equivalents (IEQ) and islet particle index (IPI) after culture. Culture time is determined by the end of the isolation until approximately 9 am on a shipping/collection day. The % recovery is calculated as 'post-isolation IEQ/pre-shipment IEQ*100.' Post-culture samples are collected for insulin and DNA as described.

*Available cell culture outcomes metadata include:* Culture time (h), Total islet equivalents after culture (IEQ), Percent IEQ recovery after culture (%), Islet particle index after culture, Insulin content after culture, Insulin:DNA ratio after culture, Insulin per IEQ after culture (ng/IEQ).

##### 3.6 Static Insulin Secretion

On the day of sample collection (see section 3.5, above) islets are handpicked to 100% purity and cultured overnight at 37°C with 5% $CO_2$. In triplicates, 15 hand-picked islets from each donor are pre-incubated at low glucose for 2 x 1 hours consecutively, and then sequentially treated with low glucose for 1 hour followed by high glucose 1 hour. Collected supernatant for low glucose, high glucose or insulin content are measured by ELISA (ALPCO STELLUX Chemiluminescence). For each donor, there are multiple glucose combinations that are used. Experimental details can be found here. Glucose pairs are:

1. 1mM → 10mM glucose
2. 1mM → 16.7mM glucose
3. 2.8mM → 16.7mM glucose

Outliers were handled in two ways. *Level 1)* A single donor-level measurement was computed as the median of the three replicate measures. Using the median instead of the mean reduced the impact of replicate-level outliers on the data. *Level 2)* The donor-level values were filtered to remove those that were more than 4 standard deviations away from the mean of all donors on a log10 scale. This removed values that were many orders of magnitude different from the population. In the Data Download tool, the data are provided at the replicate level (before outlier removal), while in-tool Omics Analysis is performed on data subjected to both Level 1 and Level 2 outlier removal.

*Available static insulin secretion metadata include:* Culture time before experiment (days), Insulin content (pg/ml), Secretion at 1 mM glucose (pg/ml), Secretion at 2.8 mM glucose (pg/ml), Secretion at 10 mM glucose (pg/ml), Secretion at 16.7 mM glucose (pg/ml), Stimulation index (1->10 mM glucose), Stimulation index (1->16.7 mM glucose), Stimulation index (2.8->16.7 mM glucose).

##### 3.7 Islet Oxygen Consumption (Seahorse assay)

At the ADI IsletCore, after an overnight culture, islets are hand-picked, and 70 islets per well are assayed in triplicate for each donor. Islets are placed in the center of the well and islet capture screens are used to keep islets in place. Plate is incubated at 37°C for 1 hour without $CO_2$, after which the plate is placed in the Xfe24 analyzer (Agilent). Basal measurements are at 2.8mM glucose, while stimulation is at 16.7mM glucose. 5uM oligomycin, 3uM FCCP and 5uM of rotenone and antimycin A are used as indicated. All solutions are made using DMEM supplemented with 1% FBS, sodium pyruvate and L-glutamine. After the run is complete, islets are collected and a DNA assay is done to normalize data. A detailed protocol is here.

*Available islet oxygen consumption (Seahorse assay) metadata include:* Basal respiration, ATP-linked respiration, Proton leak, Max glucose response, Stimulated glucose response, Max respiration, and Spare capacity.

##### 3.8 Dynamic Insulin Responses to Macronutrients

After shipment to the University of British Columbia, islets are hand picked and cultured in RPMI for 24-72hrs before the experiment to allow recovery from shipping. 65 islets are loaded per chamber and pre-incubated for 1 hour. Islets are stimulated with 6 or 15mM glucose, 5mM Leucine or 0.75mM Oleic acid/0.75mM palmitic acid as indicated. Samples are stored at -20°C until analysis with RIA. Units are presented as uU/ml. Full details are described here.

*Available dynamic insulin responses to macronutrients metadata include:* 3 mM Glucose baseline secretion (AUC), 15 mM Glucose-stimulated peak, 15 mM Glucose-stimulated secretion (AUC), 6 mM Glucose-stimulated secretion (AUC), 30 mM KCl-stimulated secretion, after glucose (AUC), 5 mM Leucine-stimulated peak secretion , 5 mM Leucine-stimulated secretion (AUC), 5 mM Leucine + 6 mM Glucose-stimulated secretion (AUC), 30 mM KCl-stimulated secretion, after leucine (AUC), 1.5 mM Oleate/palmitate-stimulated peak secretion, 1.5 mM Oleate/palmitate-stimulated secretion (AUC), 1.5 mM Oleate/palmitate _ 6 mM glucose-stimulated peak secretion, 30 mM KCl-stimulated secretion, after oleate palmitate (AUC).

##### 3.9 Single-cell Function (electrophysiology)

On the day of collection (see section 3.5 above) islets are hand-picked, dispersed to single cells and plated on 35 mm cell culture dishes as described here. 24-72 hours after plating, electrophysiology is performed by whole-cell patch-clamp using procedures and solutions outlined in publications here and here.

For earlier donors, only ‘total exocytosis’ in response to a series of 10 membrane depolarizations from -70 to 0 mV is reported. For more recent donors (from R305 onward) an expanded panel of electrophysiological measures is reported that include measures of cell size, exocytosis, voltage-dependent $Ca^{2+}$ channel activity, and voltage-dependent-$Na^{+}$ channel activity. Cell types are identified either by immunostaining (insulin/glucagon) or by single-cell RNA sequencing after the experiment. In the Omics analysis, analyses can be performed on either alpha- or beta-cells, at 1, 5 or 10 mM glucose.

> *Available single-cell function metadata include:* Size (pF), Total exocytosis (fF/pF), Early (RRP) exocytosis (fF/pF), Late exocytosis (fF/pF), Calcium entry (pC/pF), Early calcium current amplitude (pA/pF), Late calcium current amplitude (pA/pF), Sodium current amplitude (pA/pF), Half inactivation sodium current (mV).

### Ch 4 - Overview of Omics Data Types

#### 4.1 Gene expression (bulk RNA-seq)

On the day of sample collection, typically the shipment day (see section 3.5 above), 150 islets are hand picked and transferred to 1 ml of Trizol reagent. After vortexing, the sample is stored at -80°C until shipping or RNAseq at either Oxford or Stanford.

At destination, RNA is extracted from Trizol as per manufacturer instructions and concentration and quality is assessed by NanoDrop. RNA integrity is further checked by Bioanalyzer (RNA 6000 chips) and 20 uL of RNA samples at 20 ng/uL minimum is submitted to for library prep and sequencing to Novogene, on NovaSeq 6000 PE150 illumina platform, at 20M paired end reads per sample (6 G of raw data). After quality control via FastqQC (v0.11.9), the raw reads were aligned to the human genome reference (GRCh38) using STAR (v2.7.9a) with the gene annotation downloaded from the ENSEMBL database (v110). Gene expression levels were counted using featureCounts (v2.0.3). Raw counts were normalized using the TPM method to adjust sequencing depth.

Ensembl gene IDs were mapped to Entrez IDs and counts from any duplicated Entrez IDs were summed. Transcripts with over 80% zeros were filtered out. The RNAseq dataset contains seven batches, and significant batch effects were observed. Batch effects were corrected for using the Combat-Seq function in the sva R package (version 3.44.0), which preserves the count nature of the data. Counts were converted to log-counts-per-million (logCPM) using the relative log expression (RLE) method in the edgeR R package (version 3.38.4). Transcripts with logCPM variance in the lowest 20th percentile were filtered out. Since the sequencing depth was different across different batches, a substantial number of transcripts had zero counts in some batches and low but robust counts in other batches. These batch-related drop-out patterns led to large numbers of significant differentially expressed features that disappeared if individual batches were held out. To address this problem, counts with a value of zero or one were replaced with NAs, excluding these individual values from all downstream statistical analyses and leading to more stable differential expression analysis results.

The raw RNA sequencing data has been deposited in the European Genome-phenome Archive (EGA) under study number EGAS00001007241.

#### 4.2 Gene expression (Nanostring)

On the day of sample collection (see section 3.5, above) 50 islets are hand-picked at the ADI IsletCore, washed with PBS, then lysed in 100ul of RNeasy Lysis Buffer (RLT, Qiagen) containing 1% beta-mercaptoethanol. Samples are stored at -80ºC before running on Nanostring gene expression assay at BC Children Hospital (Vancouver). The assay was updated partway through data collection, where a small number of genes were removed and replaced with others, resulting in two different gene codesets. Briefly, 1ul cell lysate is used for hybridization with customized gene code sets that target 142 pancreatic islet genes and 6 reference genes. Hybridized complex is then loaded on cartridge and Nanostring nCounter to measure the target gene expression. The raw data is analyzed with nSolver4.0 software from Nanostring, producing a table of intensity values where larger values correspond to higher gene expression values. Full details are found here.

*Data Processing:* Significant batch effects were present across codesets, and so transcripts measured in both Nanostring arrays were batch corrected using the Combat function from the *sva* R package (version 3.44.0). Gene expression intensities were normalized by sample median (each value divided by the median intensity of all genes for that donor) and log2-transformed. Given the much smaller number of transcripts compared to the RNAseq data, no abundance or variance filters were applied.

**4.3 Protein expression (proteomics)**

Human islets are shipped to the University of British Columbia, hand-picked and cultured in RPMI for 24-72hrs before the experiment to allow the islets to recover from shipping. After recovery, 300 islets are hand-picked, washed with PBS and the pellet is snap frozen and stored at -80°C. LC-MS/MS analysis is performed with n = 3 technical replicates, using a NanoElute UHPLC system (Bruker Daltonics) with Aurora Series Gen2 (CSI) analytical column coupled to a Trapped Ion Mobility – Time of Flight mass spectrometer (timsTOF Pro; Bruker Daltonics, Germany) operated in Data-Independent Acquisition - Parallel Accumulation- Serial Fragmentation (DIA-PASEF) mode. Details of lysis and analysis are found here.

Protein abundances were filtered to remove any that had more than 50% missing values, median normalized, and log2 transformed. Missing values were imputed with the missForest method (random forest-based) using the *imp4p* R package (version 1.2.0). Uniprot IDs were converted to Entrez IDs, and rows with duplicate IDs were aggregated by taking the mean. Proteins with variance in the lowest 20th percentile were filtered out.

Raw proteomics data has been deposited to ProteomExchange via MASSive under dataset number PXD045422.

**4.4 Gene expression (pseudobulk RNA-seq)**

The single-cell counts from patch-seq data (section 3.5, below) were summarized at the donor by separating counts by cell type (alpha and beta). Only cells from donors with at least five cells were retained. Pseudobulk profiles were created by summing counts from all cells derived from the same donor. Ensembl gene IDs were mapped to Entrez IDs and counts from any duplicated Entrez IDs were summed. Each matrix was filtered to remove transcripts with greater than 80% zeros. Counts were converted to logCPM values using the TMM method from the edgeR R package (version 3.38.4). Due to the large variations in sequencing depth across batches, individual transcripts that had zero or one count were replaced with NA values.

**4.5 Single-cell gene expression (patch-seq)**

The collection and analysis of pancreas patch-seq data is described here and here. Reads from FASTQ files were trimmed using TrimGalore version 0.6.5 with auto-detection of adapter sequence, then aligned and counted using STAR version 2.7.9a to the GRCh38 human genome (Ensembl release 104). Cell count matrices were loaded into R and a Seurat object was generated using the Seurat R package (version 4.3.0). Only cells with patch-clamp data were included; data for additional cells are available from the listed accession numbers below. Cell types were identified by projection, using the MapQuery command from the R package Seurat, onto a large set of human islet single cell RNAseq data generated from publically available data (see below). Cells were filtered to remove any that were not alpha or beta cells, had fewer than 500 total counts across all transcripts, had fewer than 300 unique transcripts with non-zero count values, or had greater than 20% mitochondrial DNA. Ensembl gene IDs were mapped to Entrez IDs and counts from any duplicated Entrez IDs were summed. Counts were separated by cell type (alpha and beta) and glucose concentration (1, 5, and 10mM), resulting in six different matrices. Each matrix was filtered to remove transcripts with greater than 80% zeros. Counts were converted to logCPM values using the trimmed mean of M-values (TMM) method from the edgeR R package (version 3.38.4). TMM has been shown to perform better for single-cell RNAseq data compared to the RLE method because it is robust to the high drop-out rate. Individual transcripts that had zero counts were replaced with NA.

Raw sequencing reads are available in the NCBI Gene Expression Omnibus (GEO) and Sequence Read Archive (SRA) under accession numbers GSE124742 and GSE164875 and at PancDB (https://hpap.pmacs.upenn.edu/). Some unpublished data remain to be deposited upon publication.

**4.6 Single-cell RNA-seq**

Reads from FASTQ files were trimmed using TrimGalore version 0.6.5 with auto-detection of adapter sequence, then aligned to the GRCh38 human genome, Ensembl release 104, and counted using STAR version 2.7.9a except for files from GSE84133, which were aligned and counted using the inDrops protocol; files from GSE81076 and GSE85241, which were aligned and counted using the R package scruff (version 1.10.1) with Rsubread (version 1.22.2); and files from GSE269204, which were aligned and counted with Cell Ranger (10X Genomics Cell Ranger v6.1.2). Cell count matrices were loaded into R and objects generated using the R package Seurat (version 4.3.0), restricted to transcripts detected in more than 10 cells and cells with between 200 and 12,500 unique transcripts with non-zero count values; cells with percent mitochondrial transcripts in the top 25% for each dataset were also dropped. Transcript names were maintained as Ensembl identifiers to preserve the greatest number of unique transcripts. After combining all donors, transcripts expressed in fewer than 0.1% of cells were filtered. Donor HPAP-027 was dropped from the data set because there were very few cells (32 cells). The reciprocal PCA integration with references (HPAP-037, HPAP-040; HPAP-045, HPAP-050, HPAP-052, and HPAP-072) workflow from the R package Seurat (version 4.3.0) was followed, with k.anchor = 5, then FindNeighbors and FindCluster (resolution = 1.2). Cell types were first manually annotated based on the unambiguous, high level expression of marker genes in distinct clusters in UMAP space, except cytotoxic T cells, which did not form a distinct cluster but had restricted expression of *TRAC*. The remaining ambiguous cells were annotated using the MapQuery function by projecting unknown cells onto the manually annotated cells. Although these data include cells from donors with type 1 diabetes and type 2 diabetes, only cells from donors with no diabetes (ND) are visualised in the web tool for cell-type localization of called features, and are not downloadable .

Data are from the NCBI Gene Expression Omnibus (GEO) and Sequence Read Archive (SRA) under accession numbers GSE81076, GSE81547, GSE81608, GSE83139, GSE84133, GSE85241, GSE86469, GSE154126; from the European Molecular Biology Laboratory European Biology Institute (EMBL-EBI) under accession number E-MTAB-5061; from the Human Pancreas Analysis Program at PancDB up to donor HPAP-109, and from unpublished data (GSE269204).

### Ch 5 - Overview of statistical analysis methods

##### 5.1 Multiple linear regression

Linear models are used to compute omics feature-metadata associations for the bulk RNAseq, Nanostring, pseudobulk RNAseq, and bulk proteomics datasets. A linear model containing the selected metadata variable of interest and all covariates (if any) is fit to the expression or abundance levels of each feature in the selected omics dataset, using the limma R package (version 3.52.4). If the user selects a categorical metadata of interest, two dropdowns automatically appear so that a specific contrast can be specified (ie. 'type 2 diabetes' versus 'no diabetes'). Then, the coefficient of the metadata variable (continuous) or metadata contrast (categorical) of interest and its associated p-value are extracted from the model for each feature. P-values are adjusted using the false discovery rate method. The rationale for using the original limma method is that it has been shown to perform well for many omics types as long as the data are appropriately normalized and transformed. While more customized methods developed for specific omics types may perform slightly better when applied to the appropriate data type, the advantage of our approach is that it can be applied to many omics types without modification, making the results more comparable to each other and making the HumanIslets tool easier to maintain. A detailed explanation of how analyze omics data with multiple linear regression, including how to specify linear models for different research questions and how to interpret the results, is available in Protocol #2 here.

**5.5 Deconvolution analysis**

After normalization for tissue sample size, the intensity of a marker protein that is only abundant in one cell type and that has relatively constant levels is proportional to the abundance of that cell type. We assume an average of 0.55 beta cells, 0.30 alpha cells, 0.10 delta cells, and 0.05 gamma cells and use this to estimate the cell type composition of each donor using proteomics data. We computed the proportion of each cell type for each donor, according to marker genes. The overall proportion of each cell type was calculated as the median of proportion estimates across all marker genes for that cell type. Finally, the proportion of non-endocrine cells was computed as the remainder after adding the estimated proportion of beta, alpha, delta, and gamma cells and normalizing the endocrine proportions to satisfy an assumed maximum purity of 98%. When computing associations between individual omics features and cell type proportions, endocrine cell proportions were normalized as a proportion of the total endocrine signal and the non-endocrine proportion was included as a covariate to better distinguish cell type-specific signals.

Overall, this can be thought of as estimating the strength of the signal associated with four endocrine cell types, anchoring the median signal strength with prior known cell type proportions, and attributing any missing signal to trapped non-endocrine cells. It relies on the assumptions that marker genes are only expressed in one cell type and that their levels are constant across donors. Neither of these assumptions are completely satisfied, however the method should be a robust estimator so long as the majority of marker genes have heavily biased expression in one cell type and if all marker genes are not regulated in the same direction (ie. all beta cell marker genes up-regulated in donors with diabetes).

**5.6 Sources of gene annotations & link to code repository**

Entrez IDs, official gene symbols, and short gene descriptions were accessed from NCBI's FTP site on February 2, 2023 (link here), and curated for use throughout the HumanIslets tool. Copies of all associated R scripts, including functions called by the HumanIslets tool, scripts used for data processing and analysis, and scripts for generating manuscript figures are available at this repository.
